# Experimental wildflower additions increase pollination in urban agroecosystems

**DOI:** 10.1101/2025.01.23.633830

**Authors:** Emilie E. Ellis, Jill L. Edmondson, Stuart A. Campbell

**Affiliations:** School of Biosciences, The University of Sheffield, Western Bank, Sheffield, S10 2TN, U.K; Research Centre for Ecological Change, Department of Organismal and Evolutionary Biology, University of Helsinki, Finland

**Keywords:** ‘Urban pollinators’, ‘Experimental flower additions’, ‘Pollination’, ‘Bees’, ‘Moths’, ‘Hoverflies’, ‘Pollination experiment’, ‘Pollinator conservation’

## Abstract

The addition of nectar-rich flower patches in human-modified ecosystems is a common practice to mitigate pollinator declines and boost pollination. However, the benefits of these additions for pollinator communities and pollination services are rarely tested, especially in urban environments.
In a city-scale experiment we added floral resources to urban allotments and monitored the effects on bees, hoverflies and moths, and tested for improved seed set in a model crop (tomato, *Solanum lycopersicum)*.
The addition of wildflowers did not benefit all insect communities. Only social bee abundance (*Bombus* and *Apis*) benefitted from increasing floral resource area whereas other insect taxa showed no changes in abundance potentially due to the divergence in foraging patterns of moths, hoverflies, social bees and solitary bees. The addition of wildflower patches enhanced pollination by supporting a 25.3% increase in tomato seed set, providing evidence that wildflower interventions can improve urban pollination. Seed set was higher in more urban sites, suggesting an “oasis effect” where pollinating insects are concentrated into limited greenspaces. This highlights the precarity of pollination services in highly urban areas.
Our results suggest that the practice of planting wildflower patches can positively affect pollination services in urban areas. The continued promotion of flower patch addition is likely to benefit some key insect taxa, however, the common wildflower species in seed mixes may not benefit hoverflies and moths compared to bees. The taxon-specific foraging patterns we observed should inform the design and development of pollinator-friendly wildflower seed mixes.

**Societal impact statement:** Enhancing urban greenspaces for pollinator communities by planting wildflower patches is increasingly common, but their efficacy for different groups of insects (bees, hoverflies and moths) is unclear. Our city-scale experiment demonstrated that wildflower patches benefitted the abundance of social bees, but did not increase other groups. Wildflower addition increased pollination services, with an increase in seed-set in our model crop, particularly in more urban areas. Wildflower patches do benefit pollinator communities and in turn humans, through the pollination services, but the species mix may not benefit some insect groups. This suggests there is potential to improve the benefit provided by wildflower patches by redesigning pollinator-friendly seed mixes.

## Introduction

Pollination by insects is a crucial ecosystem service essential to terrestrial ecosystem function (Ollerton et al., 2011) and underpins 33% of global crop production (Potts et al., 2010). There are growing concerns for the resilience of this pollination service due to global insect pollinator declines (Potts et al., 2010; Powney et al. 2019; Wagner et al., 2021a). In an effort to halt the declines in the abundance and diversity of key pollinator groups such as bees, there has been a surge in pollinator conservation schemes, driven by policy led changes (Hall et al., 2017) and NGOs, including direct conservation measures and public engagement (e.g. ‘No-mow may’ (Plant Life, https://www.plantlife.org.uk/)) and ‘Gardening for Wildlife’ (RSPB, https://www.rspb.org.uk/)). These schemes may play an important role in supporting insect communities, making it important to understand their effects on pollination of both wild plants and crops.

Floral resources (nectar and pollen) are vital for pollinator populations (Frankie and Thorp, 2009), and there is strong evidence that floral resource availability affects the abundance and diversity of wild bee populations (Kennedy et al., 2013). Supplementing floral resources has therefore become a focus of pollinator conservation efforts, particularly in human-modified landscapes (Bommarco et al., 2013; Braman and Griffin, 2022). These measures can improve pollination services. For example, in conventional agricultural systems, wildflower planting in field margins and the restoration of hedgerows can offer abundant foraging resources and nesting sites for insect pollinators, and in some cases increase the yield of nearby insect-pollinated crops (e.g. Morandin and Kremen, 2013). The impact of floral additions on pollinators has been well-studied in agricultural contexts where the impact is generally positive (Haaland et al., 2010; but see Delphia et al., 2022), however, despite the rapid expansion of urban areas and a corresponding rise in urban agriculture, fewer studies have been conducted in cities, where the impact of floral additions at different scales remain unclear.

The expansion of cities through the process of urbanisation has been shown to be a key driver of pollinator biodiversity declines (Wagner et al., 2021a), and there are important opportunities for insect conservation through the provision of floral resources within urban greenspaces. Within cities, there is an emerging willingness of the public to participate in conservation interventions. This engagement has been attributed to overwhelming evidence of the positive relationship between well-being and immersion in nature (Russell et al., 2013), the mental and physical health benefits of gardening (Gulyas et al., 2024), and the popular media coverage of the rapid decline in pollinating insects (e.g. ‘Insect Armageddon’, The Guardian 2017). Public awareness of the benefits of insects, including pollinators, has facilitated establishment of wildflower verges and pocket parks (e.g. the All-Ireland Pollinator Plan, https://pollinators.ie/), with the assumption that these interventions carry benefits for both insect wildlife and human activity. However, empirical evidence for these benefits is still sparse and is crucial for providing informed management advice and optimising the effects of these interventions on urban wildlife.

Compared to agricultural land, greenspaces within cities can contain a high diversity of plants that are generally beneficial to pollinating insects (Clarke and Jenerette, 2015; Baldock et al., 2015). However, they are often interspersed in a matrix of impervious surfaces and other unsuitable habitat (McKinney, 2008), which limits the availability of these vital resources. Our understanding of how pollinating insects respond to surrounding urbanisation is limited due to the few comparative studies that examine the relative responses of both bee and non-bee pollinators to urbanisation, especially insects that rely on non-floral resources for the completion of their life cycles as larvae, such as hoverflies (Diptera: Syrphidae, Bates et al., 2014; Baldock et al., 2015) and moths (Lepidoptera, Ellis et al., 2023). As a result, the links between urbanisation, insect community composition and pollination services in cities remain unclear.

The effects of resource availability on pollinator diversity, particularly non-bee pollinators, could have important implications for urban pollination services. Moths complement diurnal pollination networks and account for up to one-third of plant-pollinator interactions in urban plant-pollinator networks (Ellis et al., 2023) and have complex pollen-transport networks in agricultural ecosystems (Walton et al., 2020, MacGregor et al., 2019, Alison et al., 2022). Hoverflies are diurnal pollinators with divergent life-history traits from bees and have been shown to pollinate crops and wild plants, while also contributing to biocontrol of pest species (Jauker et al., 2012; Dunn et al., 2020; Rader et al., 2020). Due to their non-floral resource or habitat requirements, it is likely that the addition of floral resources alone may not have the same benefits shown for bees (Moquet et al., 2018). However, despite their relative vulnerability to urbanisation (Baldock et al., 2015, Theodorou et al. 2019), these taxa are rarely assessed when examining the benefits of habitat restoration or floral resource supplementation. Consequently, it remains unclear whether wildflower additions represent an optimal conservation practice that supports all pollinating insects.

Within cities, horticultural spaces such as allotments present unique opportunities to evaluate the benefits of floral resource supplementation along gradients of urbanisation. Allotment sites are large greenspaces with individual plots of land (∼250 m^2^) rented by individuals or households for growing fruits and vegetables (Dobson et al., 2020) which are dependent on insect pollination. These spaces support urban agriculture and contribute to social capital and mental well-being, extending benefits beyond the plot holders to their broader communities (Dobson et al., 2021; Gulyas et al., 2024). They are excellent spaces for evaluating civic engagement with pollinator conservation (Siegner et al., 2020). The increasing demand for allotments since the COVID-19 pandemic (Lin et al., 2021; Gulyas et al., 2024) underscores the substantial opportunities to enhance their ecological and social value. Moreover, the nutritional value of these spaces is significant, for example, Gulyas et al., (2021) have shown that the UK fruit and vegetable production in allotments can supply more than half of the vegetables and 20% of the fruit consumed annually by growers, with participants achieving an average of 6.3 portions of their recommended 5-a-day, which is 70% higher than the UK national average. This highlights that allotments not only play a role in fostering food production but also in promoting healthier diets and contributing to food security. Given the crucial role of pollination in maximising many crop yields, the enhancement of pollinator habitats within allotments could further amplify their contribution to food production, thus reinforcing their value as key assets in urban food systems.

In addition to their societal benefits, allotments are among the most species-rich urban green spaces, boasting high plant (Borysiak et al., 2017) and insect (Baldock et al., 2019) diversity. Their wide distribution across urban areas provides an opportunity to simultaneously assess the effects of floral resource additions on pollinator diversity and crop production. Recent research has shown that pollen-transport networks of Lepidoptera and bees in urban allotments are disrupted by the densification of surrounding impervious surfaces (Ellis et al., 2023; Herrmann et al., 2023). However, little is known about the consequences for crop production or whether resource supplementation could mitigate urbanisation’s negative effects on different insect groups. Supplementing floral resources could increase crop yield through at least two mechanisms: 1) by increasing the number/diversity of visitors (e.g. Garibaldi et al., 2013) and/or 2) by increasing foraging intensity and subsequent pollination efficiency (e.g. Blaauw et al., 2014). We experimentally tested the benefits that supplemental floral additions have on the pollinator community diversity and crop production in urban allotments and examined the scale dependency of these processes. At three scales expanding from local-flower patch additions, site-level floral area and to larger landscape-scale urban intensity, we tested whether enhancing floral resources can: (i) influence bee and non-bee pollinator diversity and abundance; (ii) improve pollination services (seed set) and (iii) modulate the impacts landscape-scale urban intensity has on insect communities and pollination efficiency.

## Methods

### Study system

This study was carried out in 24 allotment sites throughout the growing season in 2020 (March-October 2020) in Leeds, England (53°47’47.33”N, 1°32’52.26”W). The experimental allotment sites were chosen in eight independent groups (experimental blocks) of three sites. To distribute the blocks evenly and capture a broad representation of ecological variation and urbanisation across the city, we strategically selected blocks to radiate outward from the city centre to the administrative boundary (as in Edmondson et al., 2016, Figure 1A).

**Figure 1:**
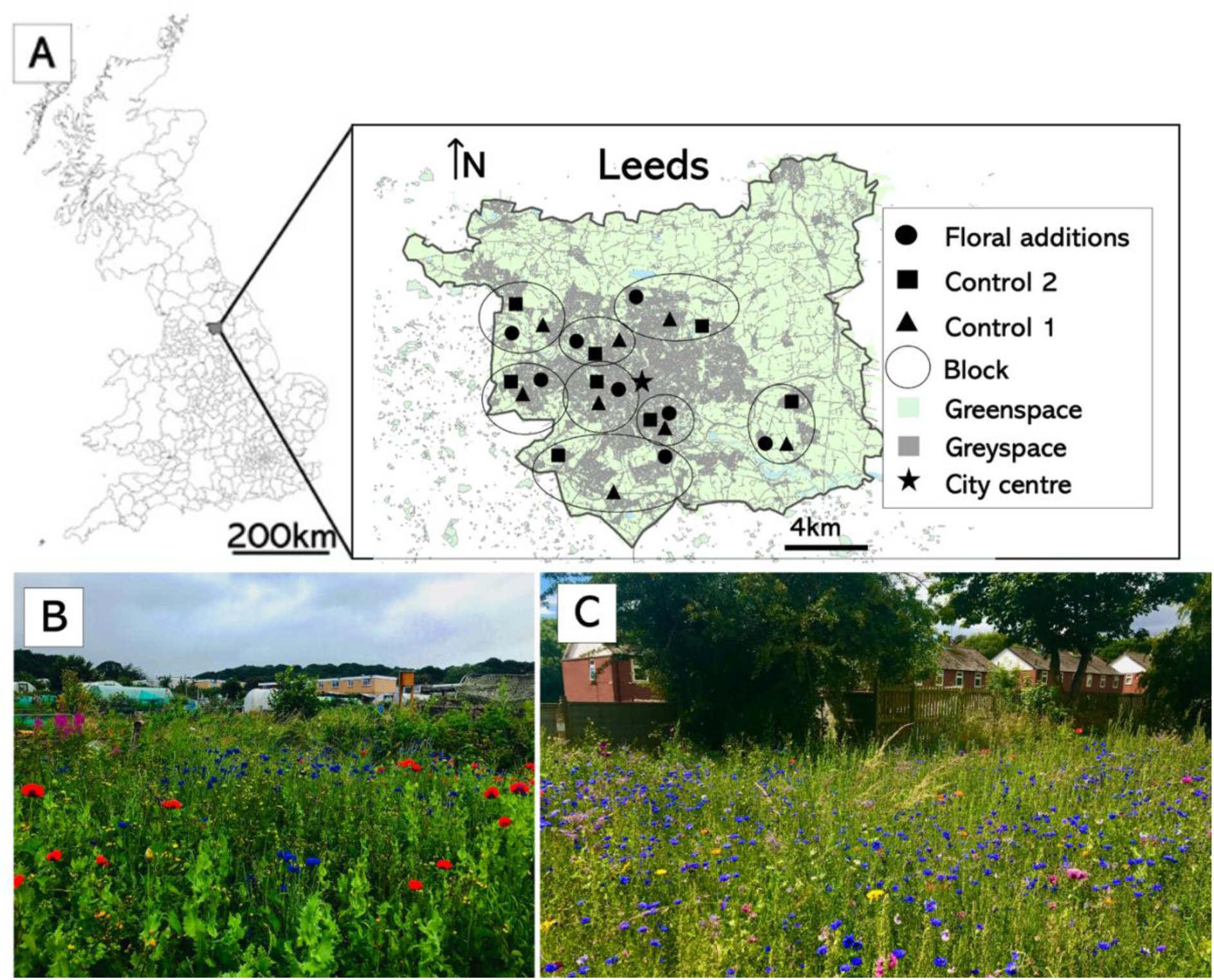
A) Left: City of Leeds’ location within the UK. Right: the site locations, treatments, and block set-up along an urbanisation gradient from the city centre. B & C) Floral additions: example of the 100m^2^ wildflower patches added in allotment sites.

### Experimental design

Each experimental block had one treatment site and two controls (control 1, control 2, Figure 2A). The treatment site was assigned a floral treatment (Figure 1). In these sites, wildflower patches (∼100 m^2^) were sown with two nectar-rich seed mixes (EuroFlor Rigby Taylor Native pollinator (biannual/perennial) and Banquet seed mix (annual/biannual); Supplementary Material Table S1; Text S1) and seven trap nests (bee hotels) were installed (Supplementary Material Figure S2). The first control site had no experimental flower patches or trap nests added. The second control site had seven trap nests installed around the site (Supplementary Material Figure S1). The uptake in all trap nests was too low (< 2%) to allow statistical analysis, but we include these sites in our analysis to control for the effect of trap nests in the wildflower addition sites. It is important to note that low uptake of bee hotels in the first year is not uncommon (McIvor and Packer, 2015).

**Figure 2:**
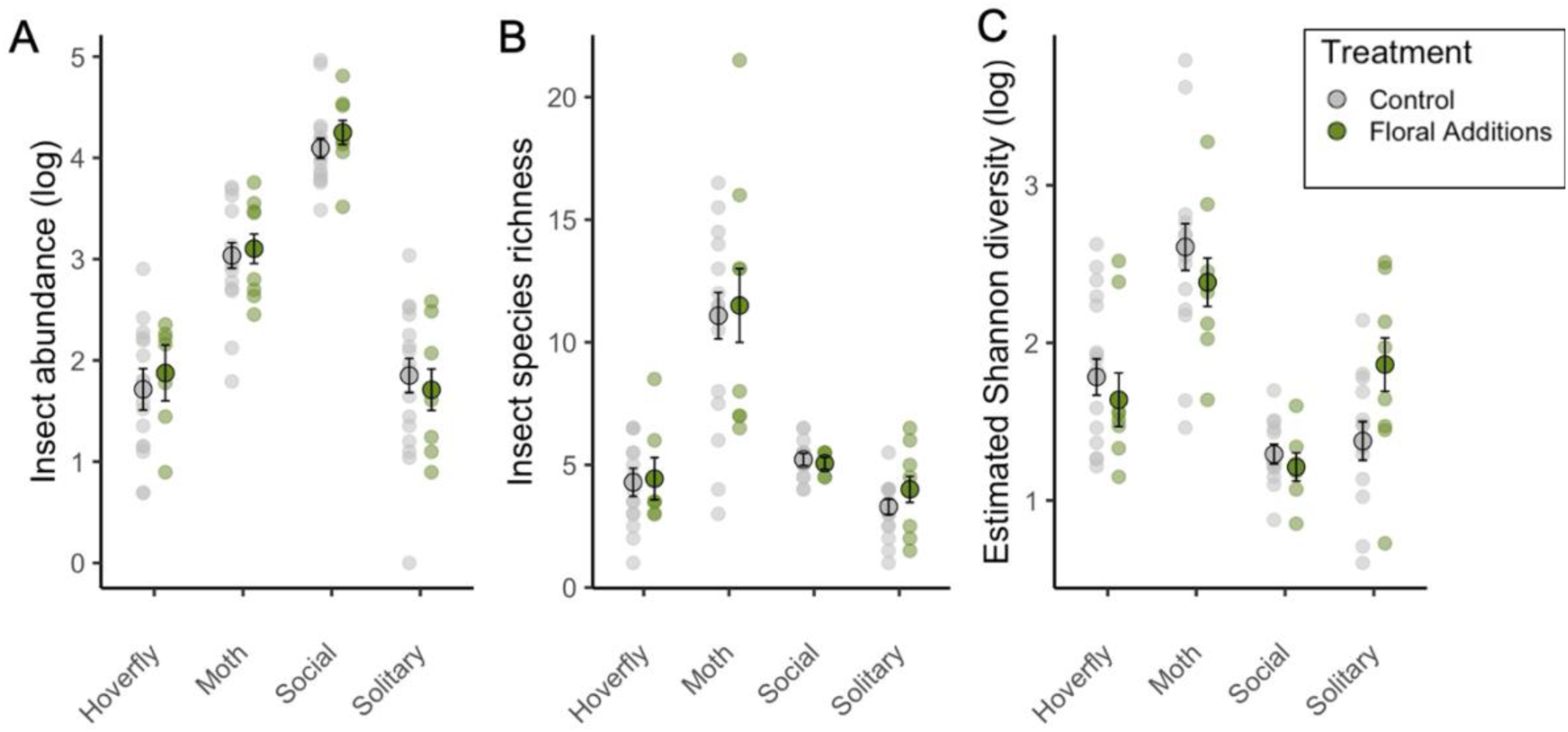
Averaged across two timepoints, showing variation in insect communities by site and taxa, comparing experimental flower patch additions (green) to control (grey) on a) abundance and b) species richness c) Estimated Shannon diversity of hoverflies, moths, and bees (social and solitary). Outlined circles denote the mean (across sites) and bars indicate standard errors.

### Sampling insects and flower-visitor interactions

To account for seasonal variation, insects were sampled on two occasions, first in early summer (20^th^ May-2^nd^ June) before the mass flowering of flower patches and again in mid- to late-summer (20^th^ July – 17^th^ August) when the flowers were in full bloom. We carried out intensive sampling at each site for each timepoint (Supplementary Material Table S2, Figure S2) and measured the species richness and abundance of three insect groups: hoverflies (Diptera: Syrphidae), bees (Hymenoptera: Anthophila) and nocturnal moths (Lepidoptera). We then compared the and visitation networks of the two diurnal insect groups. Due to the practical difficulty of observing moth-flower interactions (MacGregor et al., 2019), we did not measure moth visitation. Non-syrphid flies, despite being important pollinators (e.g. Tiusanen et al., 2016), were not included due to the limitations of our taxonomic skills.

To ensure that we captured a full representation of the communities and the site pollination efficiency, we used a diversity of sampling methods. Passive sampling included pan traps, light traps and phytometer pollination plants. We also used active sampling (walking transects and static focal surveys) to gather information about what flowers pollinators were visiting. All of which are explained in turn below, and more details are included in Supplementary Material Table S2, Figure S2.

### Diurnal pollinators (hoverflies and bees)

#### Pan traps

To sample diurnal pollinators (bees and hoverflies) we randomly placed six sets of blue, yellow and white pan traps (diameter: 7cm, height: 6cm) around each site. Each pan trap was two-thirds filled with unscented soapy water and left for five days at each of the two sampling timepoints.

#### Transects and focal surveys

To record site-level insect visitation networks, on warm calm days (between 9am-5pm) one twenty-minute transect with a sweep net was carried out through the central path of each allotment site at each timepoint and all insect-plant visitation (any event of an insect landing on a flower) were recorded. To test if the visitation patterns of insects were different in our manipulated flower patch addition sites compared to control sites, at each timepoint three ten-minute focal surveys were carried out on 0.5 x 0.5 m flower patches in both control sites and in the treatment sites. In the control sites, the focal surveys were conducted in three random flower patches around the allotment. In our treatment sites, at the first timepoint (before our experimental flower patches were flowering), these focal surveys were conducted in random flowering patches around the allotment (as in the controls). In our second timepoint, when the patches were in full bloom, the three focal surveys were all carried out in different areas of the experimental flower patch.

For both transect and focal surveys all insects and the plants they visited were recorded to the lowest taxonomic rank and aggregated by site at each timepoint. Insects were identified on the wing where possible, and when field identification was not possible, they were collected to be identified in the lab. Bees were grouped into life history categories of social and solitary following Bees Wasps & Ants Recording Society (BWARs) comprehensive life-history information (https://www.bwars.com/).

### Nocturnal pollinators (moths)

Nocturnal moths were sampled in each site during each sampling timepoint on calm, warm nights using a 12-volt portable Heath Trap (NHBS product code SK22) equipped with a 15W actinic bulb.

To ensure the sites within the blocks were closely related, we carried out each sampling method in each block specific site within three days of each other at each timepoint. The order in which blocks were sampled in the first timepoint was randomised and this randomisation was repeated at the second timepoint, as was the order of the three sites visited within the block, to minimise any temporal biases.

### Local and landscape variables

To understand the scale at which habitat variation was influencing pollinator communities and pollination, at each allotment site, a series of site-level and landscape-scale variables were estimated. We used a proxy for floral resources to estimate flowers available in each site. We mapped the area of ‘cultivated flowers’ (i.e., area of managed flower beds, flowering fruit and vegetable crops on each plot) in each site in July 2020 using visual surveys. These maps were then digitised in ImageJ (Schneider et al., 2012) and the area (m^2^) of cultivated flowers was extracted. In addition, a proxy measure for urbanisation measured at a landscape-scale was estimated for each site, by quantifying the area of impervious space in a 250m buffer surrounding each allotment using ‘Manmade’ land cover data from OS Mastermap in a geographic information system (ArcGIS version 10.7.1). We found that area of impervious surfaces and distance from city centre were negatively correlated (R^2^= 0.37, p = 0.0017, Supplementary Material Figure 3A) and across all sites, we captured a range of 0.05-0.25km^2^ of impervious surface. There was some variation in the area of impervious surface within each block despite being clustered spatially (Supplementary Material Figure 3).

### Quantifying crop pollination

To quantify differences in pollination services in our treatments, we used greenhouse-raised tomato plants (*Solanum lycopersicum,* Montello-F1 bush variety) as phytometers at each site. Though tomato is predominantly pollinated by bumblebees (*Bombus spp.*), there is evidence that they also benefit from non-buzz pollinating visitors (Cooley and Vallejo-Marin 2021). Tomatoes are commercially grown as annual plants with a global annual value of USD $10.8B (Cooley and Vallejo-Marin 2021) and are also one of the most frequently grown fruits on allotments (Edmondson et al., 2020). These attributes make them an excellent model system to quantify the ecosystem service of pollination in our study system. Seeds of tomatoes were germinated and grown for one month in a controlled environment chamber before placement at our study sites. Six tomato plants, each in individual compost growbags (Tomorite Grow Bag), and with two trusses of open inflorescences, were placed in each site during the mass flowering of the experimental flower patch additions. Four growbags were randomly placed around the site and two were placed next to flower patches in our treatment sites. On each plant, one truss was bagged throughout the experiment in the field with fine net (1 mm gauze) to prevent insect visitation and assess incidental site variation in self-pollination. Once the fruit had set, three tomatoes on each of the open and bagged trusses were harvested. All seeds were counted and the average number of seeds per plant was used as a measure of the ecosystem service of pollination.

### Data analysis

All analysis was done in R version 12 (R Core Team 2022).

### Site level effects of wildflower additions

First, we tested if the addition of wildflowers increased the total area of cultivated flowers at a site level. We used linear models (car::lm) with total area of cultivated flowers (at a site-level) as a response variable and treatment (wildflower additions, control) as an explanatory variable.

### Sampling efficiency

We used the iNEXT package (Hsieh et al., 2016) to compute diversity estimates and explore our sampling effeciency and ensure our treatment and control sites did not have uneven sample saturation/completeness. Specifically, we used sample-size-based sampling curves (i.e., plot of the diversity estimates as a function of sample size). We first estimated treatment specific species richness (across taxonomic groups and timepoints) to compare our sample effeciency across treatment sites and our two control sites. We then generated Shannon diversity estimates (a Hill number which is the exponential of Shannon entropy) by using the observed sample of abundance to compute the estimate and associated 95% confidence intervales. We calculated Shannon diversity values for each taxonomic group at each site at each time point and used it as an insect community measure (response variable) in the analysis below.

### Pollinator communities

We used lmers (lme4::lmer) and linear models (car::lm) to model the effect of treatment on the insect communities (species richness, abundance, estimated Shannon diversity) while accounting for the effects of habitat (local site-level cultivated floral area and landscape-level urbanisation) co-variates and if insect taxa identity (bees, hoverflies and moths) and timepoint influences these patterns. To explore if there were taxa-specific responses to our explanatory variables, we initially included insect taxa x treatment, and insect taxa x habitat interactions, and removed them if not significant (p>0.05). Our models had the following structure:

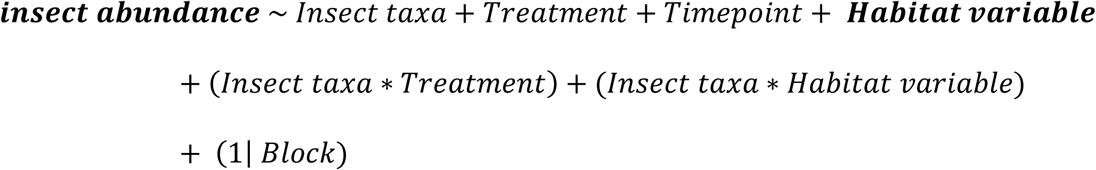

For modelling estimated Shannon diversity and species richness we used generalised linear models (car::glm) due ‘singular fit’ as there was a lack of variation of in the response variable. Thus, we fit the same model structure as above but with ‘Block’ as a main effect. For the Shannon diversity model, we also included weights which were ^1⁄^(*Upper confidence 95% − Lower confidence 95%*) of the estimated diversity measurements. We used the package emmeans (Lenth et al., 2018) to carry out any post-hoc tests when there was evidence of significant interactions (function ‘joint_test’).

### Pollination

We then tested how pollination efficiency (mean seed set per site) was influenced by our local floral additions, and site-and landscape-level variables as described above:

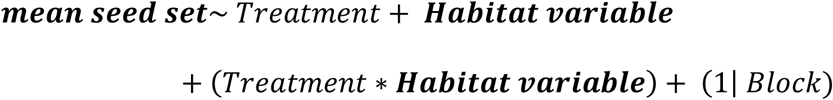

To test how bee species richness influenced seed set of tomatoes, we ran the same model as outlined above but replaced ‘Habitat variable’ with bee species richness.

### Network construction and analysis

We constructed diurnal networks based on field observations to compare the network structure metrics of hoverflies, solitary, and social bees using (bipartite::networklevel, Dormann et al., 2009). Using the data from sweep net insect-plant observations (both line-transect and focal surveys), we constructed networks for 24 sites and calculated the following network metrics: total number of plants foraged on, linkage density and host range of insect species. These metrics were used to assess how their visitation patterns were influenced by our wildflower addition treatment, the site level floral resources and the surrounding urbanisation following the model structure outlined above.

The area of cultivated flowers within the sites and the area of impervious surface were included in the models as covariates to assess how the local and landscape factors were influencing the insect communities and pollination. Exploratory analysis showed that area of impervious surface was slightly correlated with site-level area of cultivated flowers (χ² = 3.60, p= 0.058, Supplementary Material Figure S4) thus, these continuous variables were analysed in separate models.

## Results

### Fly, bee, and moth communities in urban allotments

In total, 7616 insects, belonging to 311 species were collected and observed across the two sampling periods (Table 1). Overall, we sampled between 44-53% percent of our communities with similar effort in treatment and control sites (Supplementary Material Figure S4, Table S3). The honeybee (*Apis mellifera)* accounted for 22% of the total insect community (n = 1680) and bumblebees (*Bombus spp.)* made up 27% (n = 2091). Moths were the most species rich, with 203 species recorded (2617 individuals) (Table 1; Supplementary Tables S4-S6 for full species lists). During the transects and focal collections, we recorded a total of 4611 insect-plant interactions. In total, 169 plant species were visited (Supplementary Material Table S7) and the most visited plants were *Rubus sp.* (n = 472 visits), *Origanum vulgare* (n = 368 visits), *Centaurea cyanus* (n = 302), *Jacobaea vulgaris* (n = 253), *Borago officinalis* (n = 249).

**Table 1:**
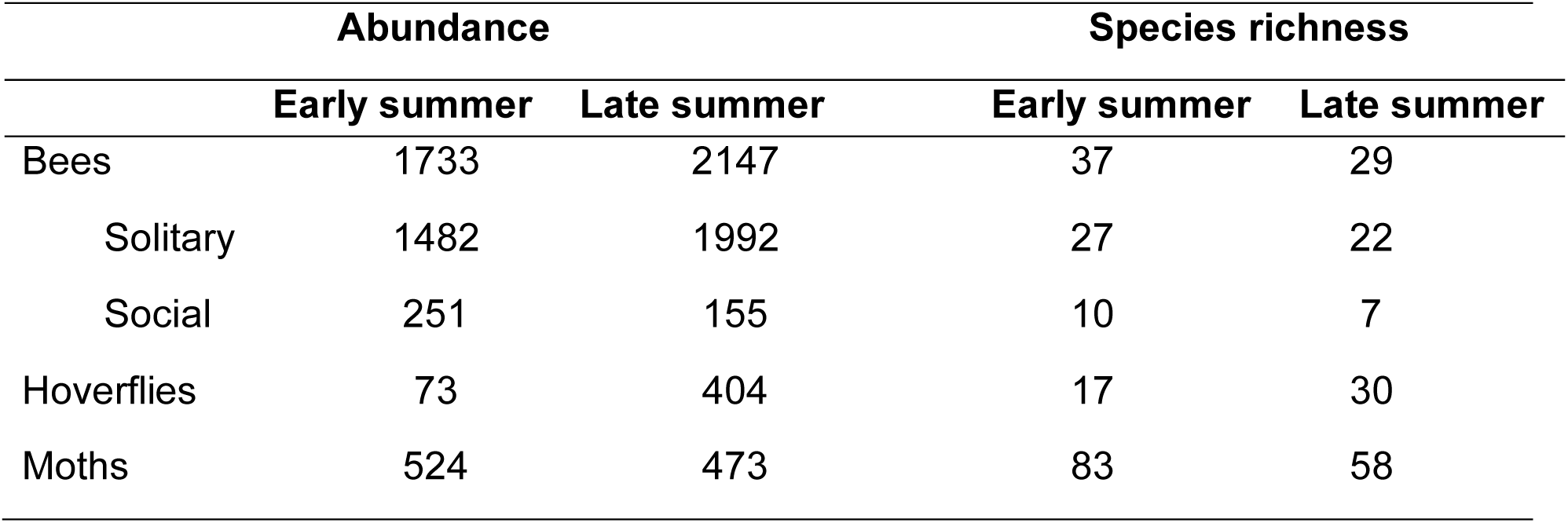
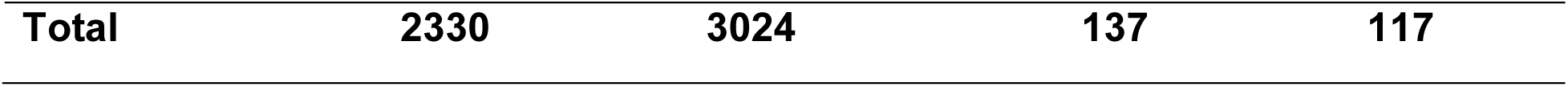
Summary of the insect species richness and abundance of bees, hoverflies and moths collected in urban agricultural sites in Leeds during the growing season 2020.

### Insect community responses to floral additions

Within our experimentally added flower patches, 47 species of plants were recorded (this was less than expected due to a drought in the spring). The most visited flowers in the patches were our sown *Centaurea cyanus* (n = 203), *Borago officinalis* (n =100), *Limnathes douglasii* (n = 56), *Sympythum officinale* (n = 51), and naturally regenerating *Cirsium vulgare* (n =48), *Jacobaea vulgaris* (n = 30) and *Sonchus oleraceus* (n = 28). There was variation among sites in non-treatment flowering area, and subsequently we found that the addition of wildflower did not significantly increase the site-level area of flowers compared to our control sites (F_(1,21)_ = 0.08, p = 0.79, Supplementary Material Figure S6). There were also no differences between the two groups of control sites (with and without bee hotels; Supplementary Material Table S8; Figure S7) and therefore justifying our discission to re-level bee nest sites into one control level in all subsequent analysis. There was also no significant difference between the insect abundance or species richness in sites with wildflower additions compared to the control (Figure 2, Supplementary Material Tables S9-S10).

Site-scale area of cultivated flowers affected insect abundance but not species richness or estimated Shannon diversity (Figure 3 A-C). Sites with higher areas of cultivated flowers had higher abundances of insects (interaction: χ² _(1,3)_ = 19.77, p = 0.0002, Supplementary Material Tables S11). This response was taxon-specific and post-hoc tests showed pattern was driven by social bees only (Figure 3B; post-hoc test: F= 23.37, p < 0.0001).

**Figure 3:**
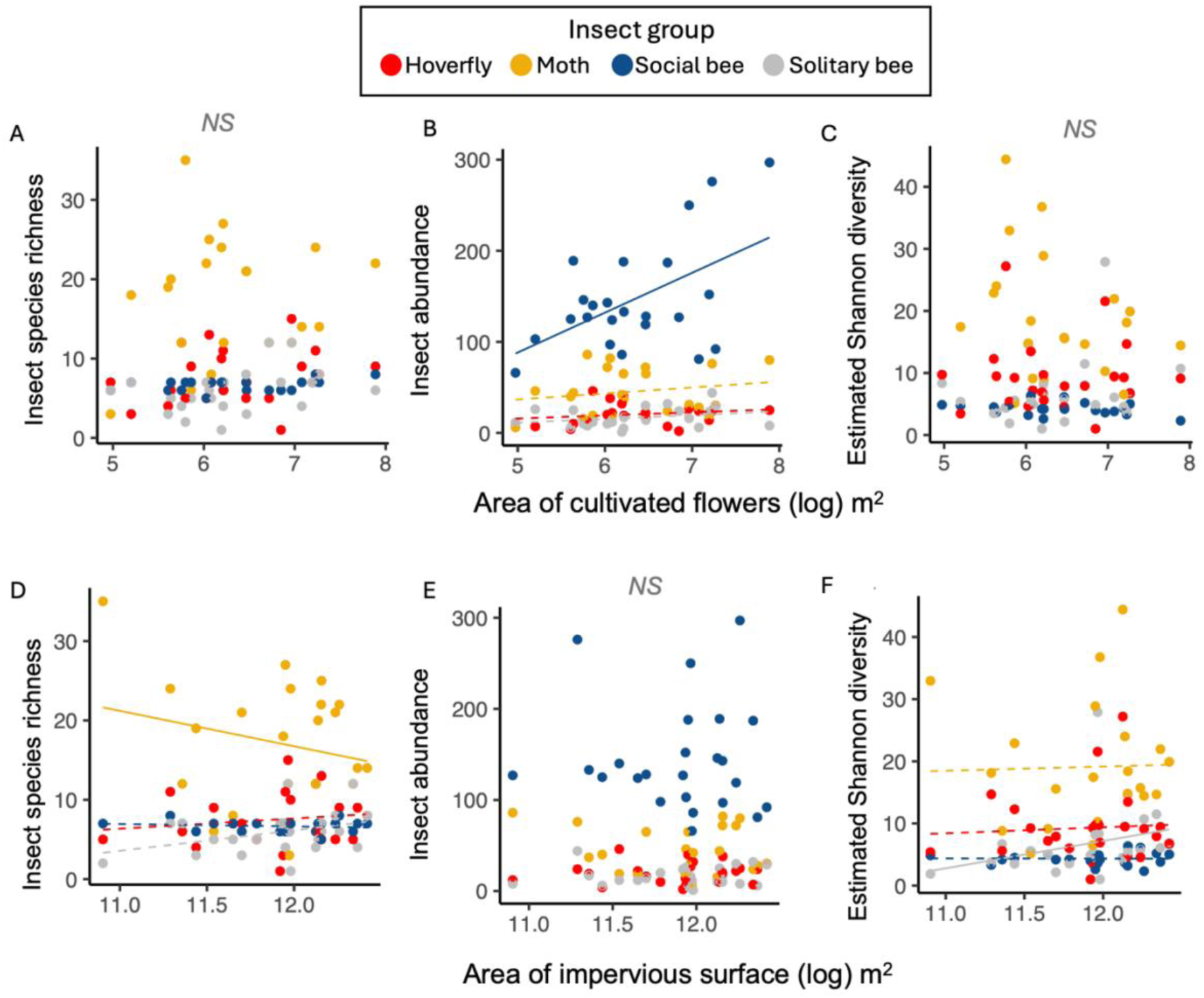
Top panels show the relationship between a) insect species richness, b) abundance and c) estimated Shannon diversity of hoverflies, moths, social bees and solitary bees with the area of cultivated flowers in allotment sites. The bottom panels show the relationship between a) insect species richness, b) abundance and c) estimated Shannon diversity of hoverflies, moths, social bees and solitary bees with the area of impervious surfaces in the 250 m surrounding the allotments sites. Points represent aggregate measures across two sampling points per site. Solid lines indicate significant linear model fits; panels with both solid and dashed lines indicate a significant interaction between insect taxa and the variable, with solid lines showing significant post-hoc groups and dashed lines non-significant groups. No lines indicate no significant effect.

There were taxon-specific responses of species richness and estimated Shannon diversity to increasing area of impervious surfaces around the site (Figure 3D and F, Supplementary Table S10, S14). Taxon-specific species richness responses to increasing impervious surface area (interaction: F_(1,3)_ = 4.56, p = 0.004) were driven by the negative response of the moth community (Figure 3D; post-hoc test: F = 10.25, p = 0.002). There were no detectable effects of urbanisation on insect abundance (Figure 3E, χ² _(1,3)_= 0.06, p= 0.80). Finally, we found some evidence that the estimated Shannon diversity significantly interacted with area of impervious surfaces (interaction: F_(1,3)_= 3.11, p = 0.03, Supplementary Material Table S14) and this was driven by the positive response of solitary bees (Figure 3F; post-hoc test: F= 0.21, p = 0.056).

### Pollination services

Pollination services were affected by a combination of local level (floral resource addition and site level cultivated flower areas) and landscape level (area of impervious surface surrounding the site) variables (Figure 4). Tomato seed set was 25.3% higher in the floral supplementation treatment compared to control and nest-only sites (F = 9.28, df = 1, p = 0.008; Figure 4A). We also found that sites with a greater proportion of surrounding impervious surface had higher seed set (F = 9.03, df = 1, p = 0.007; Figure 4B). In addition, as the area of cultivated flowers at a site level increases, so does the number of tomato seeds (F = 5.25, df = 1, p = 0.03; Figure 4C). Species richness of bees had a significant positive effect on the mean number of tomato seeds (F = 4.85, df = 1, p = 0.03; Figure 4D).

**Figure 4:**
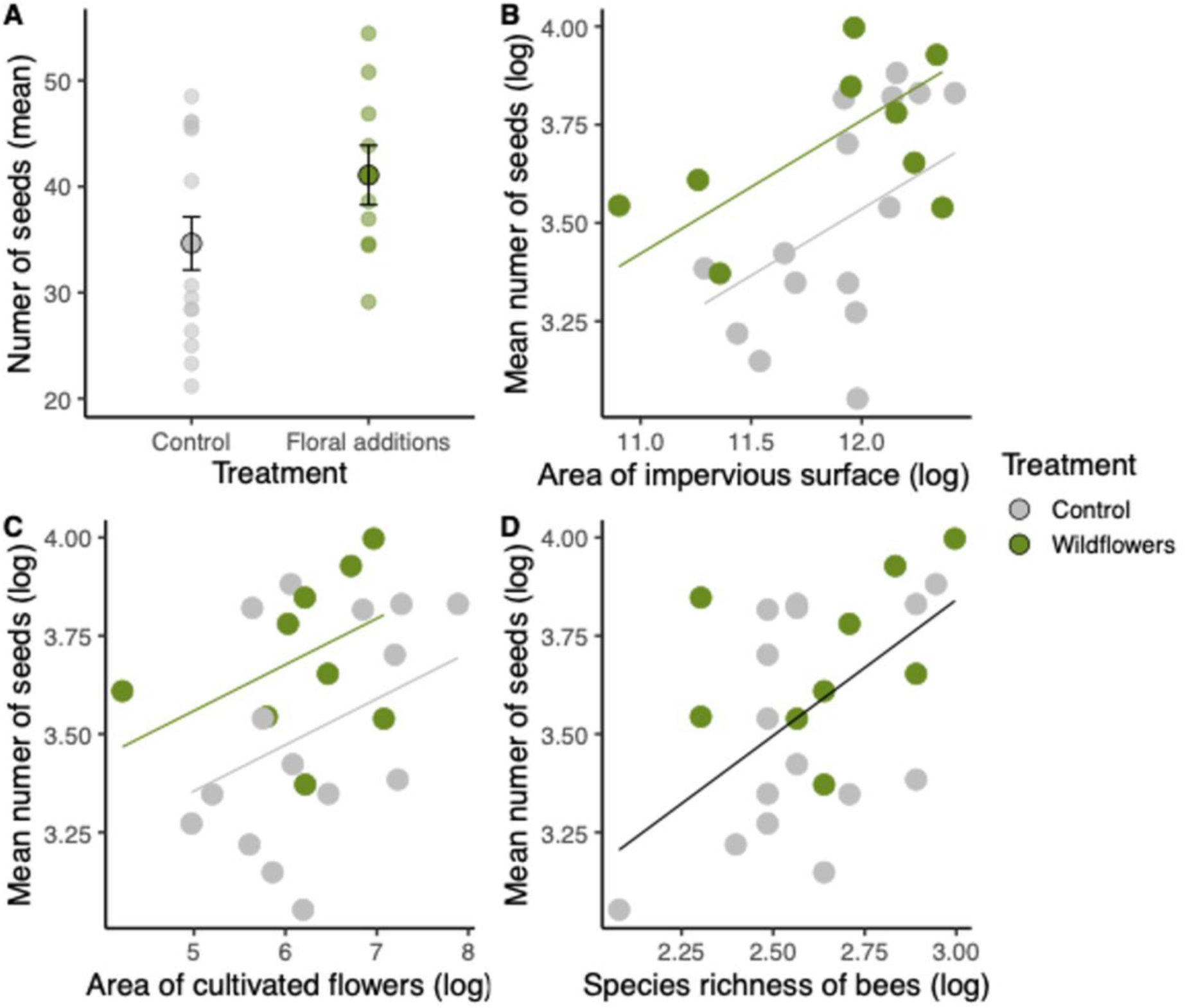
Effects of A) treatment (the addition of wildflower patches, B) urbanisation (area of impervious surface (m^2^) surrounding the sites in a 250m buffer), C) area of cultivated flowers (m^2^) in each allotment site and D) species richness of bees of tomato seed set. Data are the site observations (n = 25). A) Outlined circles denote the mean (across sites) and bars indicate standard errors. (B-C) Lines represent model fit showing significantly higher seed set in the wildflower addition treatment (green) compared to control (grey) with no significant interaction of treatment and the continuous variable. Solid black line denotes a significant overall positive effect of bee species richness on mean number of tomato seeds across all treatments (Tables S15-17).

### Visitation patterns different groups of insect pollinators

Solitary and social bees and hoverflies visited distinct floral communities (Figure 5). Only 25% of the 171 plant species recorded were visited by both bees (social, solitary) and hoverflies. The addition of hoverfly-plant interactions increased the number of plant species visited by 14%, with 23 plant species exclusively visited by hoverflies. When bees were separated based on sociality, we also found distinct plant communities visited by social (Apidae) and solitary (non-Apidae) species with only a 33% overlap (Figure 5). Social bees dominated the plant visitation observations, visiting more plant species, having higher individual host ranges and linkage density compared to solitary bees and hoverflies (Figure 5).

**Figure 5:**
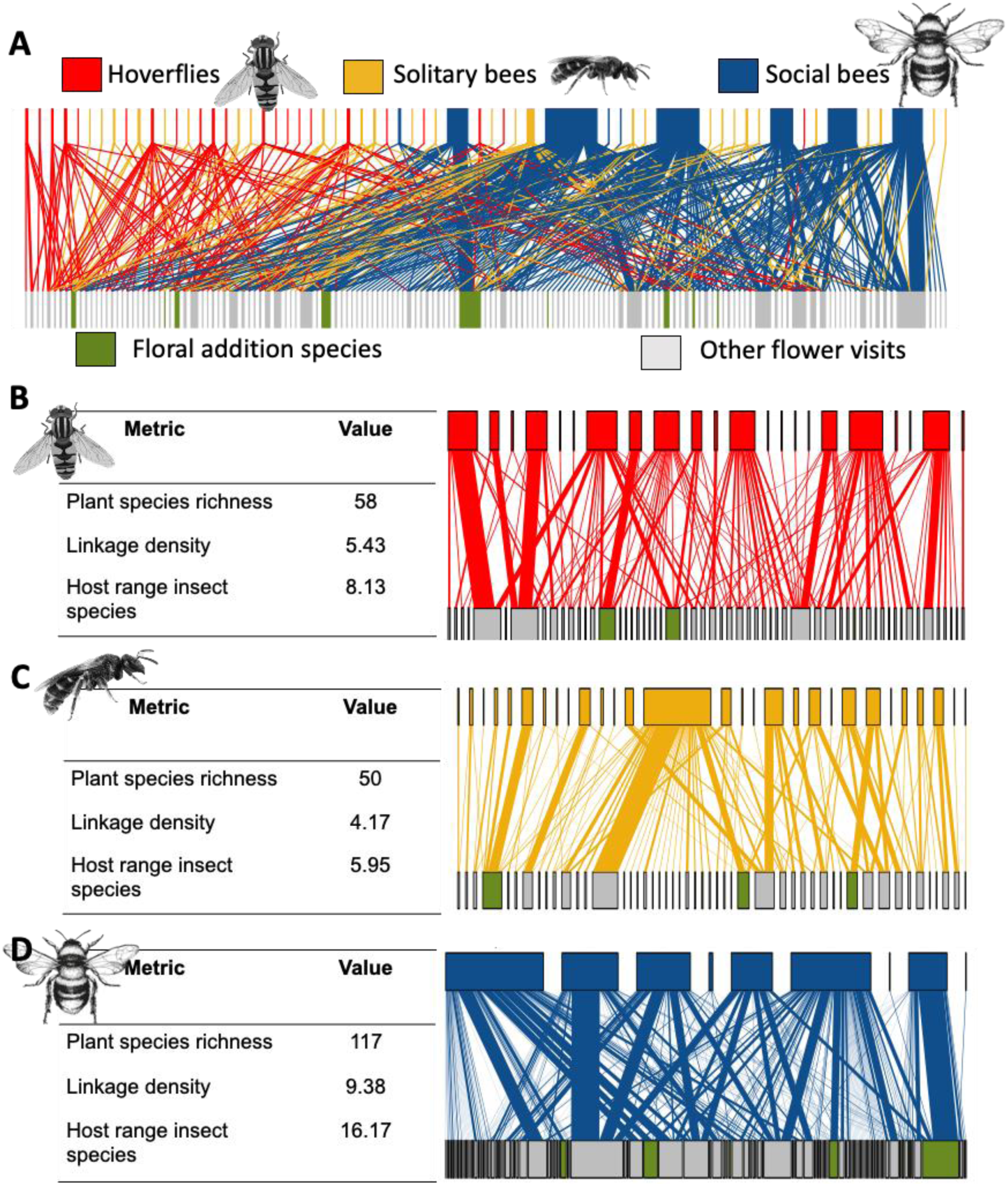
Bipartite networks showing the insect-plant visitations of diurnal pollinating insects including bees (social and solitary) and hoverflies in 24 allotment sites in Leeds, UK in summer of 2020. A) Shows all taxa in one network and illustrates the difference of visitation frequency across insect groups, the top nodes represent insect species (coloured by insect group) and lower nodes represent the plant species visited (coloured by plant species presence in the wildflower patches added to allotments) and size of node indicates number of occurrences (either insect abundance (top) or number of plant visits (bottom)). B) the hoverfly visitation network with network metrics for the illustrate network, as C) shows this for solitary bees and D) for social bees.

There was no change in pollinator-plant network structure (number of plants visited, linkage density and host range of insect) among our treatments (Supplementary Material Table S18). There were, however, significantly positive effects of the area of cultivated flowers on all diurnal network metrics tested (p < 0.05, Supplementary Material Figure S8).

## Discussion

The addition of nectar-rich flower patches in intensively modified ecosystems is a common practice to mitigate the declines of pollinators and to boost pollination services. However, despite the widespread use of wildflower planting, the benefits to insect communities and their pollination services have rarely been tested in urban areas, especially for non-bee pollinators. Using a large-scale, manipulative experiment, we found that there were complex local and landscape level drivers shaping pollinator communities and the pollination services they provide. Our experiment revealed that the addition of nectar-rich floral resources increased the pollination of our model crop by up to 25%, suggesting that the practice of planting wildflower patches can positively affect ecosystem services in urban greenspaces. However, we found that addition floral resources had no clear benefit for insect communities. Our results show that these effects were driven by complex landscape-scale interactions between urbanisation, site-level floral area and pollinator taxon, suggesting that the effective conservation of insect pollinators (especially non-bees) will require empirical testing of how resource supplementation affects specific insect groups in specific urban ecological contexts.

### Wildflower additions have strong positive effects on pollination but not insect communities

To date, the supplementation of floral resources has been predominantly tested in rural agricultural settings, where wildflower abundance can benefit the crop pollination (Albrecht et al., 2021). In an urban context, habitat amendments can have variable effects on pollination and populations of different pollinators (Griffiths-Lee et al., 2022). Of our two hypothesised mechanisms by which supplementing floral resources could increase crop yield, our results suggest that that flower additions increase foraging intensity and subsequent pollination efficiency, rather than leading to an increase in the species richness or abundance of insect communities (as in Matteson and Langellotto, 2011). This discrepancy in the impact on insect communities could be due to the attraction and redistribution of local pollinators, which then spilled over to the experimental tomato plants (Harris et al., 2023). Alternatively, local populations may be limited by non-food resources and may already be at their insect community carrying capacity for the local environment, limiting the potential for population increases (Simao et al., 2018). The improvement in pollination efficiency holds significant implications for urban agriculture, where many allotment crops rely on effective pollination to produce higher yields and quality produce (Edmondson et al., 2020). Thus, our results show that for urban growers, the addition of wildflower patches can enhance pollination and therefore food security. However, there is a need for future research to assess if these patterns are consistent for the diversity of important insect-pollinated fruit and vegetable crops.

### Taxon-specific responses to floral resources

Different taxonomic groups exhibit variation in the nectar and pollen rewards they seek (Tew et al., 2020; Matteson and Langellotto, 2011), and this may explain the variable effects the area of cultivated flowers has on insect communities, and the distinct visitation networks among insect groups (Figure 5, Ellis et al., 2023). We found distinct flower communities visited by social bees, solitary bees, and hoverflies, with only a quarter of the plant species visited by all three insect groups. Our network analysis suggest that solitary bees, hoverflies and social bees may provide complementary pollinations roles for some plant species but may be primary visitors for others. While our use of flowering plant area indicates that this can be used as a proxy for floral resource overall, future work should consider investigating the importance of within-site floral diversity (e.g., using species level vegetation mapping) in order to disentangle how site-level floral resources drive specific insect communities (as seen in McDougall et al., 2022).

Previous research has found social bees to some of the most generalist pollinating insect groups (Kleijn et al., 2015; Ollerton 2017), visiting a wide range of host plants. Consistent with this, we found social bees had the highest linkage density and diversity of flowers visited. There was also a strong positive relationship between social bees and site-level area of flowers (Figure 3B), suggesting that diet flexibility allows them to utilise the majority of floral resources, including a wider range of non-native crop species. While social insects are appreciated by the public for their pollination, our study highlights the importance of public engagement about diversity of insect pollinators beyond the charismatic social Apidae (Ollerton, 2017). Allotments may offer a real opportunity involving the public in insect conservation, as the majority of plot holder are either engaged or concerned about nature (Dobson et al., 2021) and may be able to work as citizen scientists to provide data for optimal urban greenspace management for all pollinator groups.

Finally, nocturnal insects have been shown to been highly sensitive to urbanisation compared to diurnal insects due to their sensitivity to light pollution at both larval and adult stages (Macgegor et al., 2015; Boyes et al., 2021). Consistent with this, we found evidence that moth species richness declines with urbanisation (Figure 3D). Urban moths have been shown to benefit from trees, shrubs and habitat complexity (Bates et al., 2014; Ellis and Wilkinson 2021; Ellis et al., 2023), thus, allotment cultivated flowers may not provide them with resources they require to survive. Considering recent evidence of moths being important pollinators to common pants (e.g. *Rubus* (Anderson et al., 2023), and *Trifolium* (Alison et al., 2022)). Our results highlight the need to quantify the role moths have in urban cropping systems and the functional consequences of their sensitivity to urbanisation.

### Urbanisation has complex direct and indirect effects on pollinators and pollination

Disentangling the drivers of plant-pollinator interactions as well as their pollination services is a difficult task, especially in intensely modified urban areas where there are both social and environmental factors to consider when assessing complex ecological interactions (Figure 6, Theodorou et al. 2020; McDougall et al. 2022). The counterintuitive increase in pollination in areas of greater urban intensity (Figure 4B) is potentially driven by an ecological process known as the oasis effect (Theodorou et al., 2020). Specifically, flower-rich sites (i.e. allotments) located within an inhospitable landscape (highly urban) may attract and retain insects from greater distances than sites in more floristically rich landscape (less urban), leading to a concentration of foraging, particularly by social bees, in more urban areas, which then enhances the pollination services provided. We summarise these complex drivers in a conceptual model for understanding urban pollination services (Figure 6). The strength the oasis effect is likely to also depend on variation in a range of habitat quality measures, including host plant diversity and nesting sites, that are known to differ within and among greenspace types (Baldock et al., 2019), therefore we predict that if the remaining greenspaces surrounding highly urban allotments are of high habitat quality for pollinators (e.g. gardens ad allotments), the oasis effect would be weaker as foraging becomes more dispersed (Figure 6). Future research needs to explicitly test this by incorporating the diversity of greenspace habitat types in their analysis, to understand the nuances and variability of the oasis effect.

**Figure 6:**
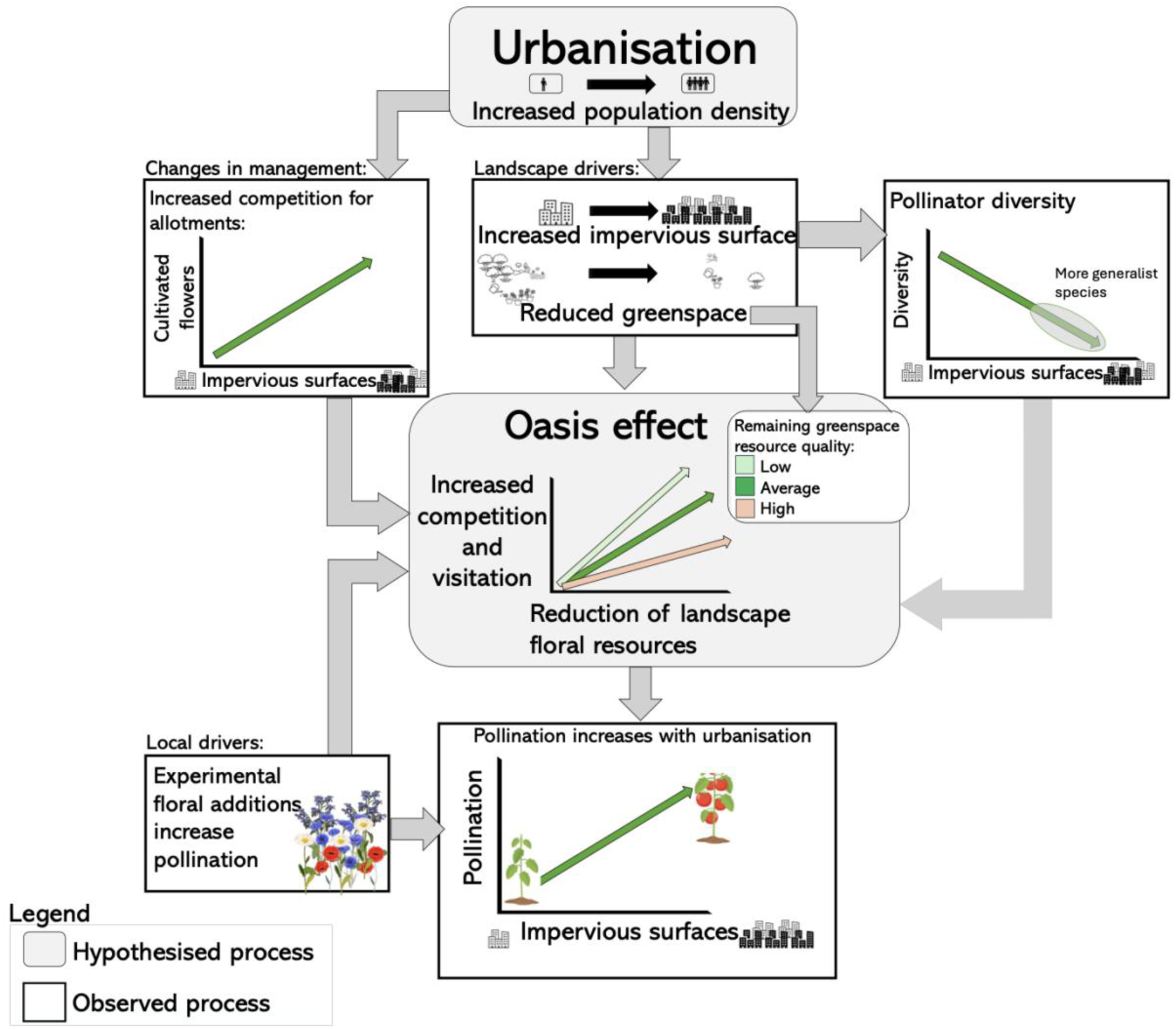
Conceptual model for the factors affecting pollination services in urban areas: Increasing urbanisation and human population density directly increase area of impervious surface (and the associated reduction of greenspace). As greenspaces area declines in highly urban areas, there are decreases in pollinator diversity and shifts to more generalist pollinator communities and increased competition and plant visitation in limited areas of floral resource. This therefore increases the pollination services within highly urban areas (oasis effect).

Increasing urbanisation was correlated with an increase in areas of cultivated flowers, suggesting that there may be local variation in social or management factors contributing to the role of allotments for pollinators. These results align with a few important studies that assess social factors in pollination conversation, (e.g. positive correlations with pollinator visits and household income, (Baldock et al., 2019). In this context, we hypothesise that one explanation could be differences in plot occupancy or turnover along our urbanisation gradient, with higher demand and turnover in plot holders leading to changes in the intensity of plot management. This positive correlation between the site-level area of cultivated flowers and urban intensity may have further increased the ’oasis effect’ of these sites for solitary bees, which were slightly more diverse in more highly urbanised sites (Figure 3F). We suggest that the robustness of these solitary bee communities will depend on changes in anthropogenic pressures that also arise from urban intensity, such as pollution and habitat degradation, and that future work should seek to explicitly link practices of the human communities using allotments with the insect communities supporting urban horticulture (Figure 6).

## Conclusion

Our study has highlighted the highly complex interactions linking pollinator communities with ecosystem services in urban environments. While our results show that the addition of wildflower patches at small scales had few direct benefits for most insect groups, they nevertheless increased the provision of pollination services, with important implications for food production in cities. The links between pollinators and seed set appear to result from complex landscape and site variation in resource concentration, coupled with taxon-specific patterns in host use (Figure 6). Finally, we show that not all insect pollinators benefited from floral resource availability, adding to a growing body of evidence suggesting that effective pollinator conservation measures need to consider the diverse resource and habitat requirements of a wider variety of pollinators across their life cycles. These results highlight the importance of understanding the foraging ecology of pollinators and the landscape ecology of urban systems in order to protect insect diversity and pollination services.

## Acknowledgments

We thank A Turner, A Maitland, B Price and E Matthews for assistance in the field and processing samples, S Falk and S Foote for help with some difficult bee and moth identifications, I Johnson for organising fieldwork consumables, and L Rogers, Leeds City Council and allotment secretaries and the other numerous allotment plot holders for site access. This study was supported by a Grantham Centre for Sustainable Futures PhD Studentship, an Engineering and Physical Sciences Research Council (EPSRC) grant to JLE (EP/N030095/1) and a Natural Environment Research Council grant to SAC (NE/R016372/1).

## Statement of authorship

EEE, SAC and JLE conceived and developed the idea. EEE and SAC set up the field experiment. EEE carried out field and lab work. EEE analysed the data. EEE wrote the first draft of the manuscript, and all authors contributed substantially to subsequent revisions.

