## Supplementary Information for "Experimental wildflower additions increase pollination in urban agroecosystems"

### **Supplementary Information for:**
### Experimental floral additions increase pollination in urban agroecosystems.

**Figure S3**: A) The relationship between the surrounding area of impervious surface (250m around the site) and the distance from the city centre (km) of each allotment site. Line indicates significant linear regression (p = 0.001). B) The average distance from the city centre of the three sites in each block, dots show the mean with error bars showing the standard error. C) The average area of impervious surface surrounding the three sites in each block, error bars denoting the standard area. .......... 6

**Table S8:** Linear mixed effect model output. Type II Analysis of Variance Table with Satterthwaite's method testing how urbanisation, area of cultivated flowers and supplement treatments (including wildflower additions, bee hotel additions and control sites), effects the abundance and species richness of bees (social and solitary), moths and hoverflies, and the seed set of tomatoes. ...........................................................................................................................24
**Table S9:** Linear mixed effect model output. Type II Analysis of Variance Table with Satterthwaite's method testing the abundance of bees (social and solitary), moths and hoverflies, and if the mean abundance differs across habitat supplement treatments, and how urbanisation influences insect abundance.......................................................................................25
**Table S10:** Linear model output. Type II Analysis of Variance Table with Satterthwaite's method testing the species richness of bees (social and solitary), moths and hoverflies, and if the mean species richness differs across habitat supplement treatments, and how urbanisation influences insect species richness, and post hoc results of significant urbanisation x insect taxa interaction. ...........................................................................................................................................26
**Table S11:** Linear mixed effect model output. Type II Analysis of Variance Table with Satterthwaite's method testing the abundance of bees (social and solitary), moths and hoverflies, and if the mean abundance differs across habitat supplement treatments, and how the area of cultivated flowers influences insect abundance. Post hoc test reported when there was a significant cultivated flowers x insect taxa interaction. ............................................................27
**Table S12:** Linear model output. Type II Analysis of Variance Table with Satterthwaite's method testing the species richness of bees (social and solitary), moths and hoverflies, and if the mean species richness differs across habitat supplement treatments, and how the area of cultivated flowers influences insect species richness. ..........................................................................28
**Table S13:** Linear model output. Type II Analysis of Variance Table with Satterthwaite's method testing the estimate Shannon diversity- derived from the abundance-based rarefaction- of bees (social and solitary), moths and hoverflies, and if the mean estimated Shannon diversity differs across habitat supplement treatments, and how the area of cultivated flowers influences insect Shannon diversity...........................................................................................................................29
**Table S14**: Linear model output. Type II Analysis of Variance Table with Satterthwaite's method testing the estimate Shannon diversity- derived from the abundance-based rarefaction- of bees (social and solitary), moths and hoverflies, and if the mean estimated Shannon diversity differs across habitat supplement treatments, and how the area of impervious surfaces effect insect Shannon diversity. Post hoc test reported when there was a significant urbanisation x insect taxa interaction............................................................................................................................30

#### Supplementary Figures:

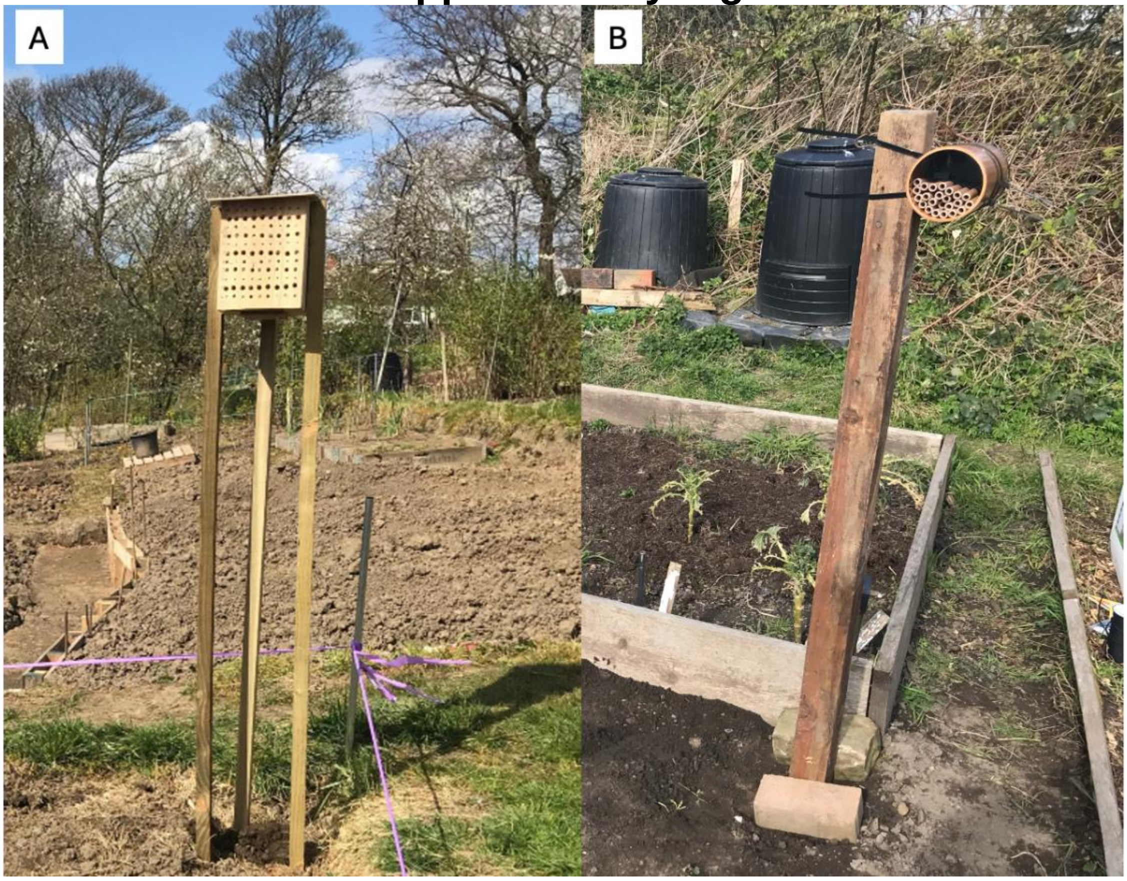

**Figure S1:** Examples of the two types of bee hotels (trap nests) used in our control sites (‘control 2’).

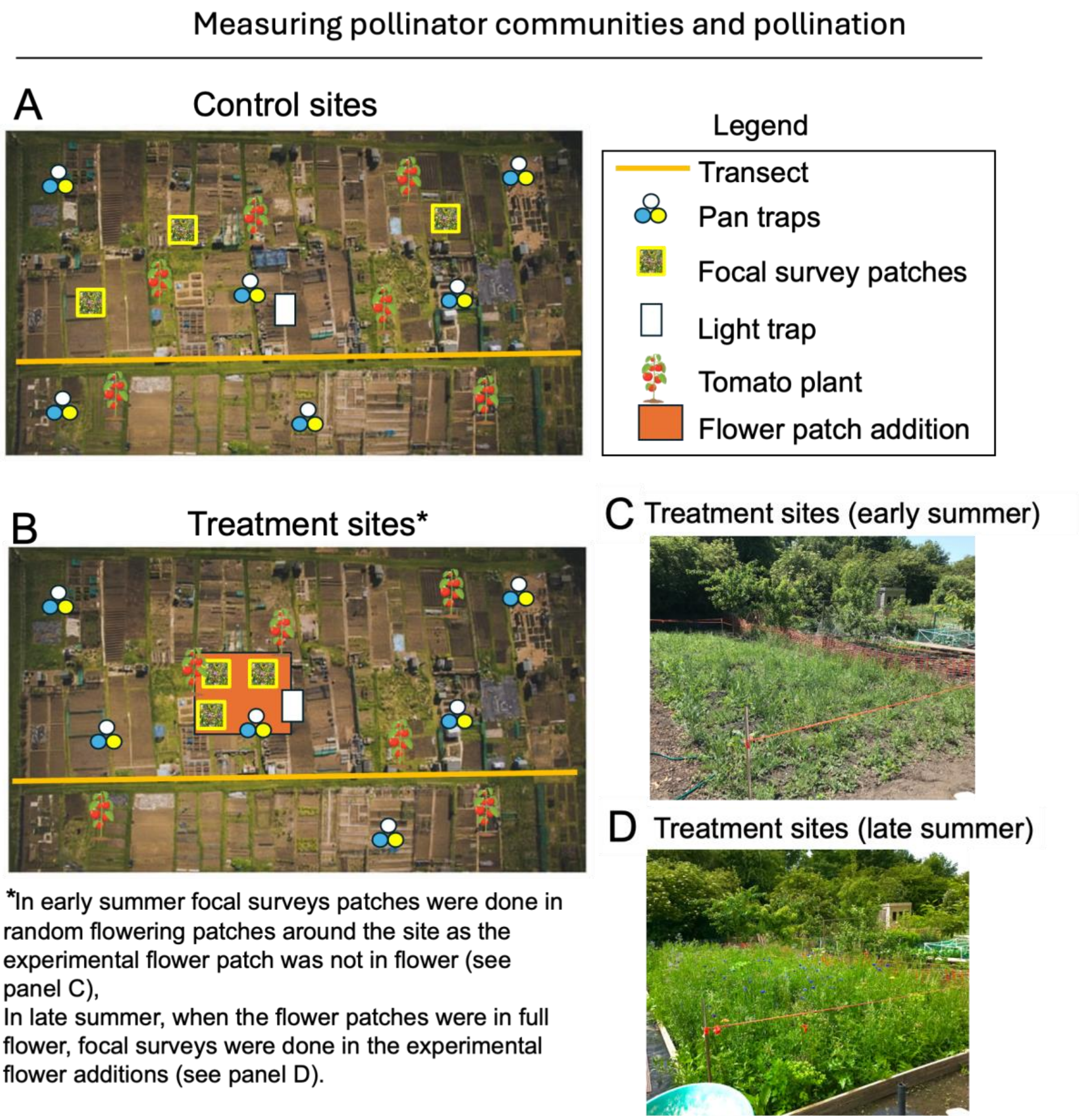

**Figure S2:** An example of the experimental set up of all sampling protocols, A) showing control site set up and B) showing treatment (where a flower patch is added) site set up. The sampling of treatments sites varied: C) in early summer when patches were not flowering sites were sampled as controls as shown in panel A, however in D) in late summer when patches were flowering, sampling was changes, as shown in panel B.

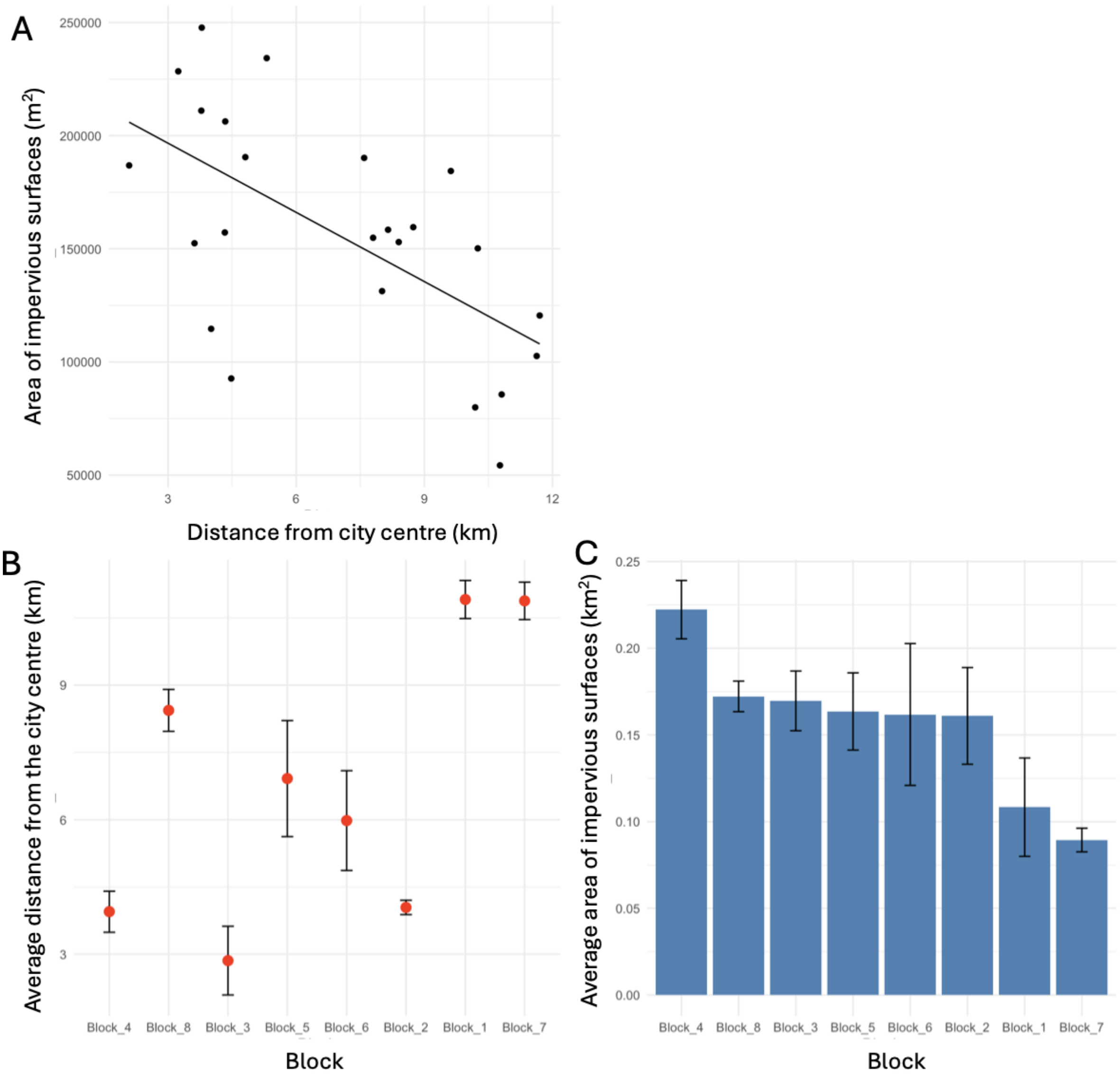

**Figure S3**: A) The relationship between the surrounding area of impervious surface (250m around the site) and the distance from the city centre (km) of each allotment site. Line indicates significant linear regression ($p = 0.001$). B) The average distance from the city centre of the three sites in each block, dots show the mean with error bars showing the standard error. C) The average area of impervious surface surrounding the three sites in each block, error bars denoting the standard area.

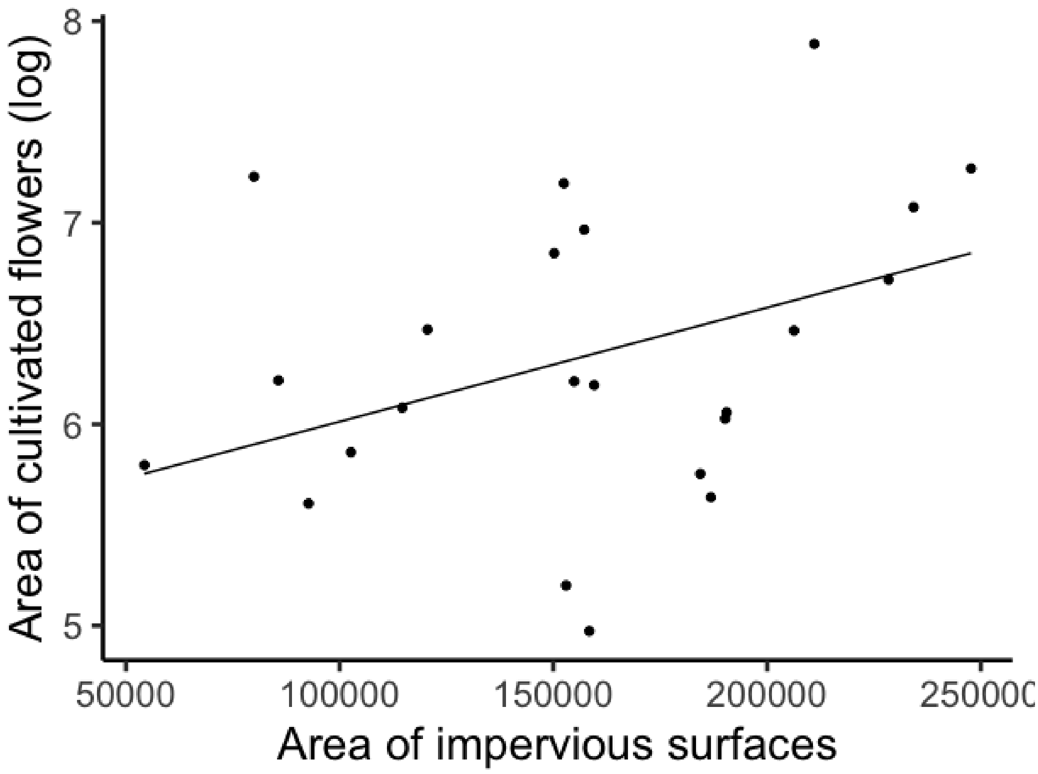

**Figure S4:** Area of cultivated flowering plants recorded during visual surveys in 24 allotment sites in Leeds increases as area of grey space surround each site ($m^2$ log) increases. Line represents linear model fitted ($\chi^2_{(1)}$= 3.60, p= 0.058).

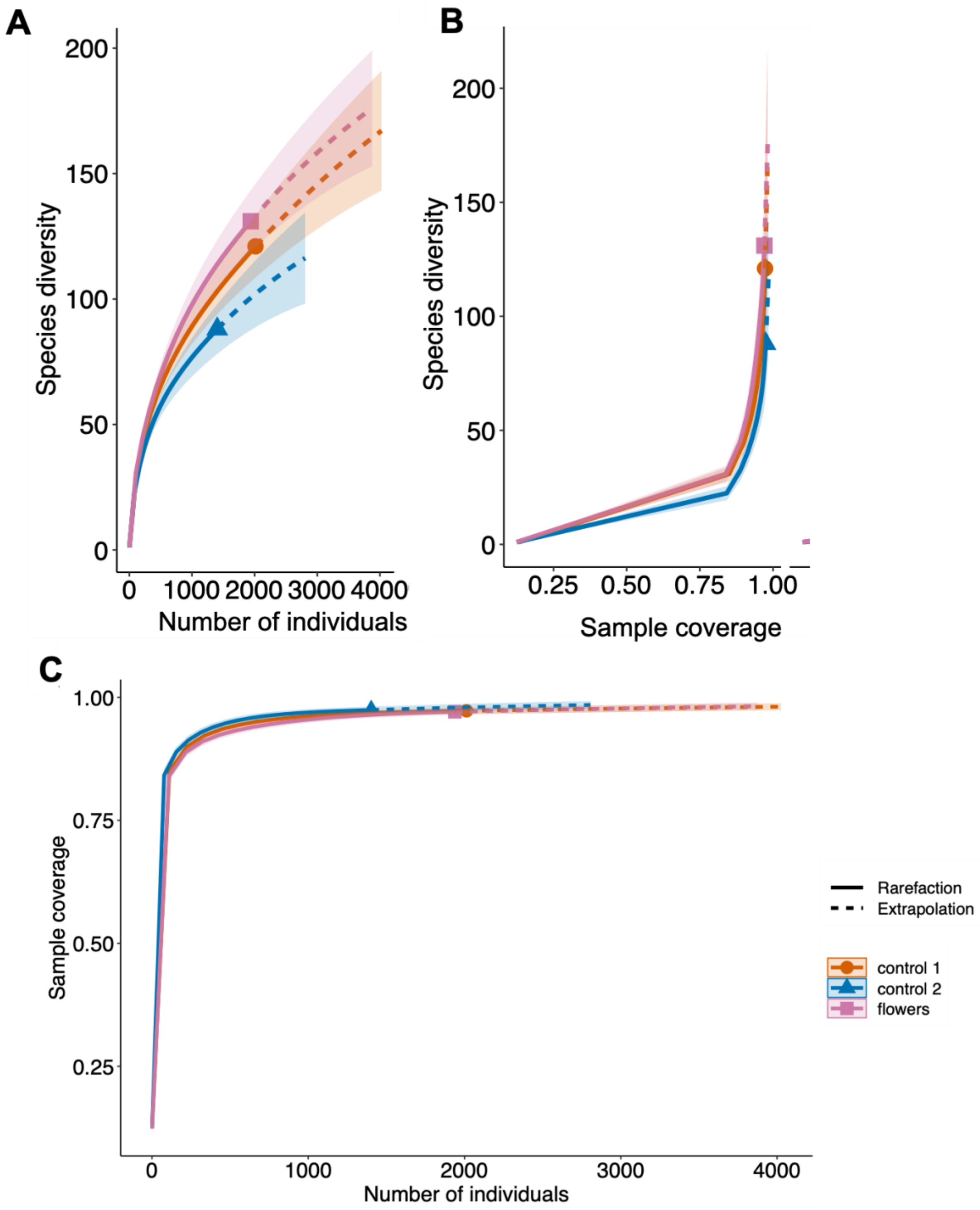

**Figure S5**: A) Sample-size-based rarefaction curves, B) coverage-based rarefaction sampling curves, C) sample completeness curve of insect pollinating communities all separated by their site's treatments (control 1, 2 and the flower patch addition). Samples rarefied by site (n=8 in each treatment).

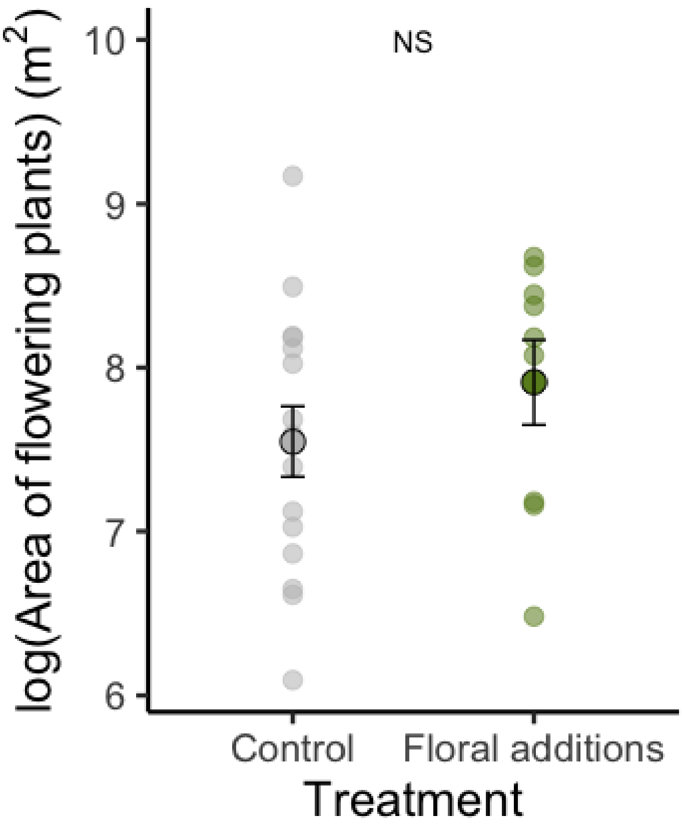

**Figure S6:** Comparing the area of flowering plants in allotments sites: Control, where no floral resources were added and Floral additions, where ~100m$^2$ of wildflowers flower were added to the sites. Light points show site total, dark point show the mean per treatment and the standard error. NS = Type II Analysis of Variance with Satterthwaite's method showed no significant different between the treatments ($F = 2.43$, $df = 1$, $p = 0.13$).

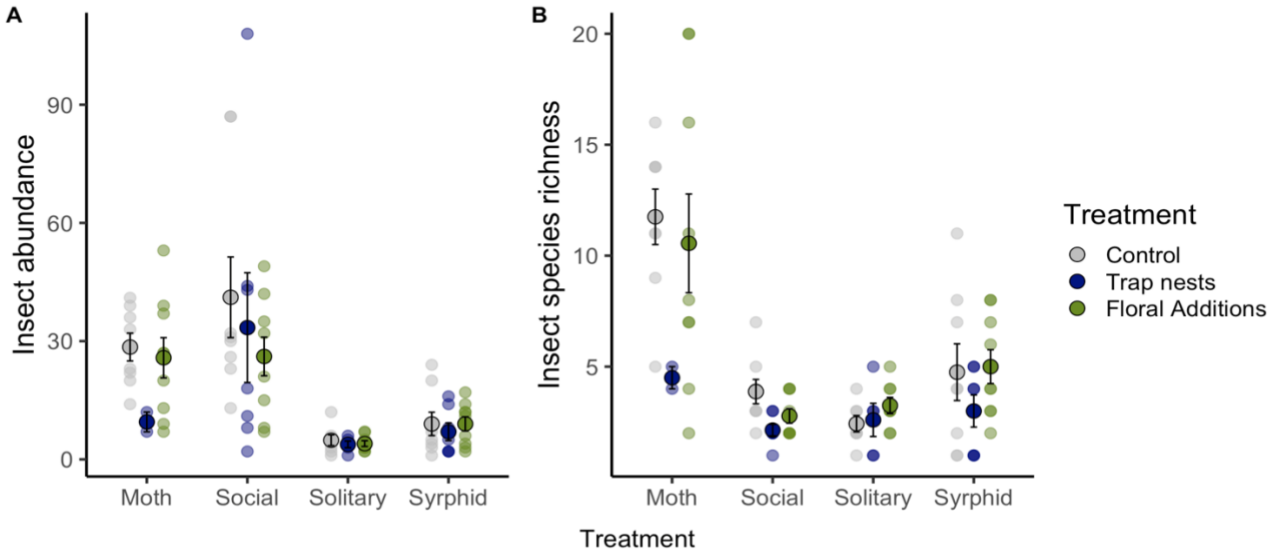

**Figure S7:** Effect of wildflower additions on a) abundance and b) species richness of hoverflies, moths, and bees (social and solitary). Colours indicate addition of wildflower treatment (green), bee hotels (navy) and control (grey) allotment sites. Outlined circles denote the mean (across sites) and bars indicate standard errors.

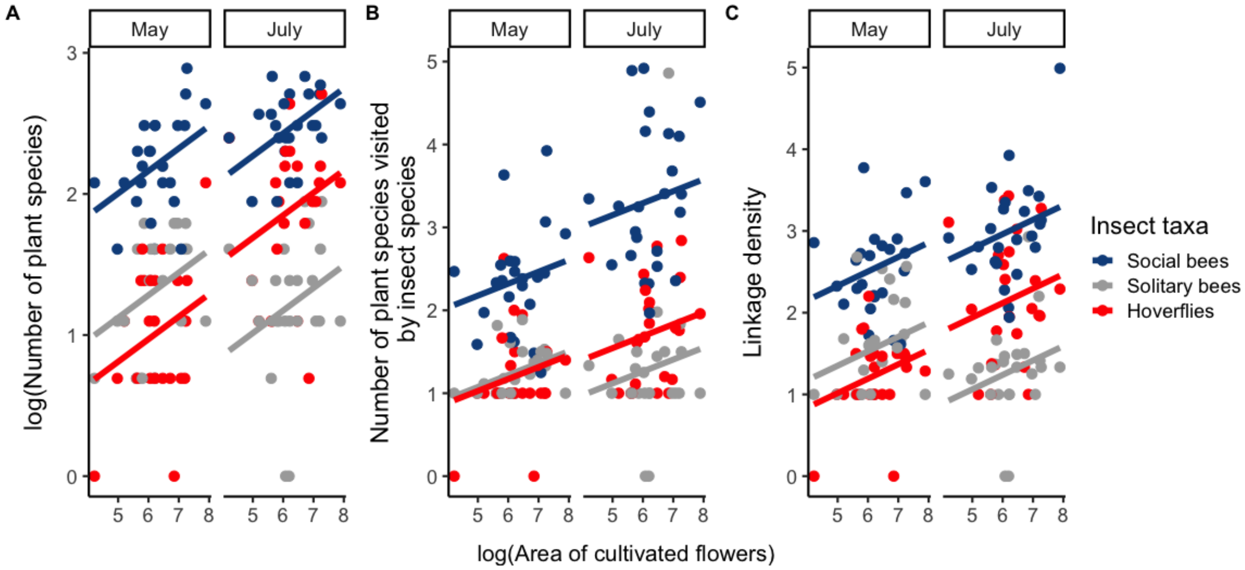

**Figure S8** Positive effects on insect visitation network metric of diurnal pollinators (hoverflies, social bees, solitary bees) as area of cultivated flowers increases in allotment sites in both early summer (May) and mid-summer (July). Lines show fitted models, with significant main effect of area of cultivated flowers ($p < 0.05$), significant insect group*time interactions ($p < 0.001$) and non-significant interaction between insect group* area cultivated flowers ($p > 0.05$).

### Supplementary Tables:

**Table S1:** Species list of seed mix of EuroFlor and Rigby Taylor Native pollinator and Banquet seed mix (for a more complete list directly contact authors).

| **Species** |
|---|
| *Lotus corniculatus* |
| *Trifolium pratense* |
| *Centaurea cyanus* |
| *Vicia sativa* |
| *Daucus carota* |
| *Papaver rhoeas* |
| *Achillea millefolium* |
| *Myosotis alpestris* |
| *Digitalis purpurea* |
| *Allium schoenoprasum* |
| *Aquilegia vulgaris* |
| *Borago officinalis* |
| *Calendula officinalis* |

**Table S2:** The details of the methods used to sample insect communities and measure pollination effeciency. Specifically indicating the target taxa, the description, the time of sampling per sampling, the number of times carried out during the sampling period and the total time of sampling across our study.

| Method | Measuring | Description | Time per site | Number per timepoint | May | July | TOTAL TIME OF SAMLING |
|---|---|---|---|---|---|---|---|
| Transect | Bees and hoverflies | Walking through the centre of the allotment site, recording any pollinator-flower interaction, identifying insects on the wing or catching them to ID in the lab by us or experts. | 20mins | 1 | ✓ | ✓ | 16 hours |
| Pan traps | Bees and hoverflies | Six sets of pan traps (1 yellow, 1 blue and 1 white) were set up around the site and emptied after 5 days. | 5 days | 6 sets in each site at each timepoint | ✓ | ✓ | 5760 hours |
| Focal surveys | Bees and hoverflies | Three ten-minute focal surveys were carried out on 0.5 x 0.5 m flower patches in each site.<br>In control sites these were randomly chosen flower patches, In the treatment sites these were also carried out in random flower patches in the first time point (before flowering) and then within the added flower patches for the second time point (during flowering) | 10mins | 3 | ✓ | ✓ | 24 hours |
| Moth traps | Moths | A light trap was set up in each site for a night at each timepoint | ~8hours | 1 | ✓ | ✓ | 384 hours |

| | | | | | | | |
|---|---|---|---|---|---|---|---|
| Phytometers | Pollination | Six tomatoes were placed around the allotment to measure pollination throughout the time series | ~2 months | 6 | ✓ | ✓ | 34, 560 hours per plant (144 plants) |

**Table S3:** Observed species richness, diversity estimates (and errors) of pollinator communities in our different treatments: controls and the addition of flower patches (8 sites in each treatment). Proportion sampled is observed divided by the estimated species richness.

| Treatment | Observed species richness | Estimator species richness | s.e. | Lower confidence interval | Lower confidence interval | Proportion sampled (%) |
|---|---|---|---|---|---|---|
| Control 1 | 121.0 | 272.1 | 48.3 | 177.6 | 366.8 | 44.5 |
| Control 2 | 88.0 | 159.9 | 23.9 | 113.0 | 206.9 | 55.0 |
| Flowers | 131.0 | 247.0 | 36.6 | 175.3 | 318.6 | 53.0 |

**Table S4:** A list of hoverfly (Diptera:Syrphidae) species recorded in 24 allotments in Leeds (U.K.) in 2020 along insect-pollinator transects and using pan traps and the number of individuals observed (total) across two collection (June and July).

| Insect species | Abundance |
|---|---|
| *Episyrphus balteatus* | 82 |
| *Syrphidae spp.* | 78 |
| *Sphaerophoria scripta* | 55 |
| *Platycheirus albimanus* | 50 |
| *Syritta pipiens* | 50 |
| *Syrphus ribesii* | 41 |
| *Syrphus vitripennis* | 33 |
| *Merodon equestris* | 24 |
| *Helophilus pendulus* | 23 |
| *Eupeodes corollae* | 19 |
| *Eupeodes luniger* | 16 |
| *Eristalis tenax* | 9 |
| *Eristalis arbustorum* | 8 |
| *Scaeva pyrastri* | 8 |
| *Melanostomas scalare* | 7 |
| *Platycheirus angustatus* | 5 |
| *Chrysotoxum festivum* | 4 |
| *Dasysyrphus albostriatus* | 4 |
| *Eumerus funeralis* | 3 |
| *Myathropa florea* | 3 |
| *Neoascia podagrica* | 3 |
| *Pipizella viduata* | 3 |
| *Platycheirus sculatus agg.* | 3 |
| *Xylota segnis* | 3 |
| *Eristalis pertinax* | 2 |
| *Platycheirus clypeatus* | 2 |
| *Volucella bombylans* | 2 |
| *Baccha elongata* | 1 |
| *Chloromyia formosa* | 1 |
| *Chrysotoxum bicinctum* | 1 |
| *Epistrophe eligans* | 1 |
| *Epistrophe grossulariae* | 1 |
| *Eristalis inticaria* | 1 |
| *Eristalis nemorum* | 1 |
| *Eupeodes latifasciatus* | 1 |
| *Heringia spp.* | 1 |
| *Heringia vitripennis* | 1 |
| *Melanostoma mellinum* | 1 |
| *Meliscaeva auricollis* | 1 |
| *Sphaerophoria indet* | 1 |
| *Spherophoria spp.* | 1 |

**Table S5:** A list of nocturnal moth (Lepidoptera) species recorded in 24 allotments in Leeds (U.K.) in 2020 using light traps (Heath) traps and the number of individuals observed (total) across two collection (June and July).

| **Insect species** | **Abundance** |
|---|---|
| *Noctua pronuba* | 326 |
| *Agrotis exclamationis* | 240 |
| *Chrysoteuchia culmella* | 218 |
| *Hoplodrina spp.* | 165 |
| *Apamea monoglypha* | 116 |
| *Oligia strigillis agg.* | 116 |
| *Mythimna impura* | 101 |
| *Lacanobia oleracea* | 60 |
| *Axylia putris* | 53 |
| *Idaea aversata* | 40 |
| *Eudonia lacustrata* | 37 |
| *Mesapamea secalis agg.* | 36 |
| *Eilema lurideola* | 30 |
| *Celypha striana* | 28 |
| *Agriphila straminella* | 22 |
| *Eupithecia sp.* | 22 |
| *Adelidae spp.* | 21 |
| *Apamea sp.* | 20 |
| *Cydia pomonella* | 19 |
| *Hedya nubiferana* | 19 |
| *Xestia triangulum* | 18 |
| *Oligia fasciuncula* | 17 |
| *Crocallis elinguaria* | 16 |
| *Hofmannophila pseudospretella* | 15 |
| *Notocelia uddmanniana* | 15 |
| *Bryophila domestica* | 14 |
| *Celphya lacunana* | 14 |
| *Campaea margaritaria* | 13 |
| *Deilephila elpenor* | 12 |
| *Xanthorhoe fluctuata* | 12 |
| *Anania hortulata* | 11 |
| *Biston butularia* | 11 |
| *Herminia tarsipennalis* | 10 |
| *Autographa gamma* | 9 |
| *Geometrid spp.* | 9 |
| *Ypsolopha dentella* | 9 |
| *Eulithis prunata* | 8 |
| *Scoparid* | 8 |
| *Acentria ephemerella* | 7 |
| *Dysstroma truncata truncata* | 7 |
| *Mythimna ferrago* | 7 |
| *Tortricidae* | 7 |
| *Tyria jacobaeae* | 7 |
| *Agapeta hamana* | 6 |
| *Aphomia sociella* | 6 |
| *Caradrina morpheus* | 6 |
| *Coleophora spp.* | 6 |
| *Ephiphyas postvittana* | 6 |
| *Eucosma cana* | 6 |
| *Geometridae* | 6 |
| *Peribatodes rhomboidaria* | 6 |
| *Cosmia trapezina* | 5 |
| *Emmelina monodactyla* | 5 |
| *Eudonia angustea* | 5 |
| *Glypjopterix spp.* | 5 |
| *Oligia strigilis agg.* | 5 |
| *Opisthograptis luteolata* | 5 |
| *Anania coronata* | 4 |
| *Diarsia mendica* | 4 |
| *Hadena compta* | 4 |
| *Mythimna pallens* | 4 |
| *Ourapteryx sambucaria* | 4 |

| | |
|---|---|
| *Phalera bucephala* | 4 |
| *Phlogophora meticulosa* | 4 |
| *Pterophorus pentadactyla* | 4 |
| *Rivula sericealis* | 4 |
| *Triodia sylvina* | 4 |
| *Xestia xanthographa* | 4 |
| *Abrostola tripartita* | 3 |
| *Aporophyla lueneburgensis* | 3 |
| *Ditula angustiorana* | 3 |
| *Helcytogramma rufescens* | 3 |
| *Hemithea aestivaria* | 3 |
| *Hydriomena furcata* | 3 |
| *Hypena proboscidalis* | 3 |
| *Lobesia abscisana* | 3 |
| *Mesoligia furuncula* | 3 |
| *Naenia typica* | 3 |
| *Noctuid spp.* | 3 |
| *Thera britannica* | 3 |
| *Abraxas grossulariata* | 2 |
| *Acleris forsskaleana* | 2 |
| *Agrotis segetum* | 2 |
| *Archips podana* | 2 |
| *Barred Yellow (Cidaria fulvata)* | 2 |
| *Blastodacna hellerella* | 2 |
| *Bryotropha sp.* | 2 |
| *Carpet moth* | 2 |
| *Clepsis consimilana* | 2 |
| *Diachrysia chrysitis* | 2 |
| *Eucosma sp.* | 2 |
| *Eudonia mercurella* | 2 |
| *Eudonia spp.* | 2 |
| *Eupithecia pulchellata pulchellata* | 2 |
| *Herminia grisealis* | 2 |
| *Lampropteryx suffumata* | 2 |
| *Lozotaenia forsterana* | 2 |
| *Noctudidae* | 2 |
| *Ochropleura plecta* | 2 |
| *Patania ruralis* | 2 |
| *Scotopteryx chenopodiata* | 2 |
| *Spilarctia luteum* | 2 |
| *Acronicta aceris* | 1 |
| *Acronicta leporina* | 1 |
| *Acronicta tridens* | 1 |
| *Agrotis puta* | 1 |
| *Alcis repandata* | 1 |
| *Amphipyra pyramidea* | 1 |
| *Anania perlucidalis* | 1 |
| *Apamea lithoxylaea* | 1 |
| *Apotomis sp.* | 1 |
| *Apterogenum ypsillon* | 1 |
| *Autographa jota* | 1 |
| *Bactra spp.* | 1 |
| *Bee moth* | 1 |
| *Blastobasis adustella* | 1 |
| *Borkhausenia fuscescens* | 1 |
| *Bryotropha terrella* | 1 |
| *Carpatolechia proximella* | 1 |
| *Celypha rufana* | 1 |
| *Cidaria fulvata* | 1 |
| *Cnephasia spp.* | 1 |
| *Coleophora albitarsella* | 1 |
| *Coleophora mayrella* | 1 |
| *Crambus perlella* | 1 |
| *Cydia splendana* | 1 |
| *Diarsia brunnea* | 1 |
| *Dichrorampha alpinana* | 1 |
| *Dioryctria schuetzeella* | 1 |

| | |
|---|---|
| *Donacaula forticella* | 1 |
| *Dysstroma spp.* | 1 |
| *Dysstroma truncata* | 1 |
| *Ecliptopera silaceata* | 1 |
| *Eilema complana* | 1 |
| *Ennominae* | 1 |
| *Epirrhoe alternata* | 1 |
| *Eucosma campoliliana* | 1 |
| *Eulithis populata* | 1 |
| *Eupithecia centaureata* | 1 |
| *Eupithecia exiguata* | 1 |
| *Eupithecia pulchellata* | 1 |
| *Eupithecia subfuscata* | 1 |
| *Euthrix potatoria* | 1 |
| *Euzophera pinguis* | 1 |
| *Evergestis forficalis* | 1 |
| *Gymnoscelis rufifasciata* | 1 |
| *Gypsonoma dealbana* | 1 |
| *Gypsonoma sociana* | 1 |
| *Habrosyne pyritoides* | 1 |
| *Hedya pruniana* | 1 |
| *Hylaea fasciaria* | 1 |
| *Iaspeyria flexula* | 1 |
| *Idaea biselata* | 1 |
| *Idaea dimidiata* | 1 |
| *Laothoe populi* | 1 |
| *Leucoma salicis* | 1 |
| *Luperina testacea* | 1 |
| *Mimas tiliae* | 1 |
| *Mormo maura* | 1 |
| *Myelois circumvoluta* | 1 |
| *Noctua comes* | 1 |
| *Noctua janthina* | 1 |
| *Noctua orbona* | 1 |
| *Nola cucullatella* | 1 |
| *Notocelia sp.* | 1 |
| *Oligia latruncula* | 1 |
| *Pandemis cerasana* | 1 |
| *Pharmacis fusconebulosa* | 1 |
| *Photedes fluxa* | 1 |
| *Phragmatobia fuliginosa* | 1 |
| *Plutella xylostella* | 1 |
| *Pseudosciaphila branderiana* | 1 |
| *Pseudotelphisa scallella* | 1 |
| *Ptilophora plumigera* | 1 |
| *Pyrausta aurata* | 1 |
| *Ryacionia buliana* | 1 |
| *Scoparia ambigualis* | 1 |
| *Scoparia ancipitella* | 1 |
| *Scopula floslactata* | 1 |
| *Scrobipalpa acuminatella* | 1 |
| *Scythropia crataegella* | 1 |
| *Smerinthus ocellatus* | 1 |
| *Spilonota ocellana* | 1 |
| *Subacronicta megacephala* | 1 |
| *Timandra comae* | 1 |
| *Tortrix spp.* | 1 |
| *Xanthorhoe decoloraria* | 1 |
| *Xestia c nigrum* | 1 |
| *Xestia ditrapezium* | 1 |
| *Yponomeutidae spp.* | 1 |

**Table S6:** A list of bee (Hymenoptera) species recorded in 24 allotments in Leeds (U.K.) in 2020 along insect-pollinator transects and using pan traps and the number of individuals observed (total) across two collection (June and July).

| Insect species | Abundance |
|---|---|
| *Apis mellifera* | 1680 |
| *Bombus terrestris agg.* | 778 |
| *Bombus pascuorum* | 458 |
| *Bombus lapidarius* | 291 |
| *Bombus pratorum* | 283 |
| *Bombus hypnorum* | 224 |
| *Hylaeus communis* | 181 |
| *Bombus hortorum* | 51 |
| *Lasioglossum calcetum* | 36 |
| *Lasioglossum leucopus* | 32 |
| *Osmia leaiana* | 31 |
| *Hylaeus hyalinatus* | 30 |
| *Lasioglossum cupromicans* | 27 |
| *Osmia bicornis* | 23 |
| *Halictus tumulorum* | 22 |
| *Colletes daviesanus* | 19 |
| *Megachile centuncularis* | 19 |
| *Osmia caerulescens* | 15 |
| *Solitary bee spp.* | 12 |
| *Andrena bicolor* | 8 |
| *Lasioglossum smeathmanellum* | 8 |
| *Megachile willughbiella* | 7 |
| *Osmia spp.* | 7 |
| *Andrena minutula* | 6 |
| *Andrena cineraria* | 5 |
| *Anthophora furcata* | 5 |
| *Halictus rubicundus* | 5 |
| *Lasioglossum spp.* | 4 |
| *Lasioglossum villosulum* | 4 |
| *Andrena nigroaenea* | 3 |
| *Anthophora plumpies* | 3 |
| *Bombus sylvestris* | 3 |
| *Lasioglossum morio* | 3 |
| *Andrena haemorrhoa* | 2 |
| *Andrena semilaevis* | 2 |
| *Lasioglossum albipes* | 2 |
| *Megachile ligniseca* | 2 |
| *Andrena albipes* | 1 |
| *Andrena fulva* | 1 |
| *Bombus jonellus* | 1 |
| *Bombus rupestris* | 1 |
| *Bombus vestalis* | 1 |
| *Coeliooxys elongata* | 1 |
| *Hylaeus signatus* | 1 |
| *Lasioglossum fratellum* | 1 |
| *Lasioglossum rufitarse* | 1 |
| *Megachile versicolor* | 1 |
| *Nomada ruuficornis* | 1 |
| *Nomada spp.* | 1 |

**Table S7:** Aggregated plant species across all sites both within flower patches and around the sites during transects and focal surveys in 24 allotments in Leeds (U.K.) and the total number of visits by bees and hoverflies during two collections (June and July 2020).

| **Species** | **Number of visits** |
|---|---|
| *Origanum vulgare* | 368 |
| *Centaurea cyanus* | 302 |
| *Jacobaea vulgaris* | 251 |
| *Borago officinalis* | 244 |
| *Symphytum officinale* | 243 |
| *Cirsium vulgare* | 240 |
| *Rubus spp. (bramble)* | 238 |
| *Rubus spp. (raspberry)* | 233 |
| *Allium schoenoprasum* | 168 |
| *Symphytum officinale* | 167 |
| *Limnanthes douglasii* | 132 |
| *Lavandula angustifolia* | 120 |
| *Ranunculus spp.* | 114 |
| *Calendula officinalis* | 113 |
| *Phacelia tanacetifolia* | 84 |
| *Rubus spp. (bramble)* | 78 |
| *Centaurea montana* | 63 |
| *Thymus vulgaris* | 63 |
| *Allium sphaerocephalon* | 57 |
| *Leucanthemum vulgare* | 55 |
| *Lapsana communis* | 44 |
| *Fragaria ananassa* | 43 |
| *Sonchus oleraceus* | 37 |
| *Trifolium repens* | 35 |
| *Nepeta spp.* | 33 |
| *Eschscholzia californica* | 32 |
| *Lupinus polyphyllus* | 32 |
| *Geranium palustre* | 29 |
| *Myosotis sylvatica* | 28 |
| *Rubus spp. (logan)* | 28 |
| *Aquilegia vulgaris* | 27 |
| *Allium nigrum* | 25 |
| *Rapistrum rugosum* | 25 |
| *Linaria purpurea* | 24 |
| *Digitalis purpurea* | 22 |
| *Epilobium ciliatum* | 22 |
| *Taraxacum spp.* | 22 |
| *Tropaeolum majus* | 22 |
| *Leucanthemum x superbum* | 20 |
| *Silene dioica* | 20 |
| *Vicia sepium* | 20 |
| *Malva arborea* | 18 |
| *Brassica oleracea* | 17 |
| *Buddleja* | 17 |
| *Dipsacus fullonum* | 17 |
| *Papaver rhoeas* | 17 |
| *Salvia rosmarinus* | 16 |
| *Allium ampeloprasum* | 15 |
| *Helenium spp.* | 15 |
| *Sinapis arvensis* | 15 |
| *Centranthus ruber* | 14 |
| *Tripleurospermum inodorum* | 14 |
| *Anthriscus sylvestris* | 13 |
| *Campanula persicifolia* | 13 |
| *Geranium pyrenaicum* | 13 |
| *Papaver somniferum* | 13 |
| *Veronica persica* | 13 |
| *Bellis perennis* | 12 |
| *Chamaenerion angustifolium* | 12 |
| *Crocosmia crocosmiiflora* | 12 |

| | |
|---|---|
| *Epilobium hirsutum* | 12 |
| *Helianthus spp.* | 12 |
| *Helichrysum italicum* | 12 |
| *Lamium purpureum* | 12 |
| *Geum urbanum* | 11 |
| *Trifolium pratense* | 11 |
| *Vicia faba* | 11 |
| *Capsella bursa-pastoris* | 10 |
| *Dahlia spp.* | 9 |
| *Geranium phaeum* | 9 |
| *Nepeta cataria* | 9 |
| *Raphanus raphanistrum* | 9 |
| *Verbena bonariensis* | 9 |
| *Veronica longifolia* | 9 |
| *Echium vulgare* | 8 |
| *Geranium endressii* | 8 |
| *Papaver alpinum* | 8 |
| *Stachys byzantina* | 8 |
| *Ammi majus* | 7 |
| *Chenopodium album* | 7 |
| *Carduus crispus* | 6 |
| *Fumariia offiicinalis* | 6 |
| *Hypericum cerastoides* | 6 |
| *Monarda punctata* | 6 |
| *Papaver orientale* | 6 |
| *Schizanthus pinnatus* | 6 |
| *Arabis blepharophylla* | 5 |
| *Calystegia sepium* | 5 |
| *Foeniculum vulgare* | 5 |
| *Iris pseudacorus* | 5 |
| *Knautia arvensis* | 5 |
| *Persicaria maculos* | 5 |
| *Phaseolus coccineus* | 5 |
| *Rosa canina* | 5 |
| *Allium giganteum* | 4 |
| *Anemone tomentosa* | 4 |
| *Antirrhinum majus* | 4 |
| *Brassica rapa* | 4 |
| *Coriandrum sativum* | 4 |
| *Dietes iridioides* | 4 |
| *Geranium himalayense* | 4 |
| *Glebionis coronaria* | 4 |
| *Polemonium caeruleum* | 4 |
| *Rosa rugosa* | 4 |
| *Inula helenium* | 3 |
| *Lamium hybridum* | 3 |
| *Myosotis arvensis* | 3 |
| *Penstemon campanulatus* | 3 |
| *Rudbeckia hirta* | 3 |
| *Senecio vulgaris* | 3 |
| *Weigela florida* | 3 |
| *Centaurea nigra* | 2 |
| *Convolvulus arvensis* | 2 |
| *Dimorphotheca pluvialis* | 2 |
| *Diplotaxis tenuifolia* | 2 |
| *Epilobium montanum* | 2 |
| *Eucosma sp.* | 2 |
| *Fumaria capreolata* | 2 |
| *Geranium macrorrhizum* | 2 |
| *Graptophyllum pictum* | 2 |
| *Gypsophila spp.* | 2 |
| *Hypericum perforatum* | 2 |
| *Ligustrum ovalifolium* | 2 |
| *LoniceraÂ caprifolium* | 2 |
| *Oenothera glazioviana* | 2 |
| *Polygonum aviculare* | 2 |
| *Psilostrophe cooperi* | 2 |

| | |
|---|---|
| *Rosa californica* | 2 |
| *Rosa spp.* | 2 |
| *Solanum tuberosum* | 2 |
| *Spirea douglasii* | 2 |
| *Stachys sylvatica* | 2 |
| *Thlaspi arvense* | 2 |
| *Veronica officinalis* | 2 |
| *Achillea millefolium* | 1 |
| *Allium cepa* | 1 |
| *Amsinckia menziesii* | 1 |
| *Anthemis cotula* | 1 |
| *Clarkia amoena* | 1 |
| *Daucus carota* | 1 |
| *Dianthus barbatus* | 1 |
| *Euphorbia helioscopia* | 1 |
| *Fuchsia magellanica* | 1 |
| *Heuchera micrantha* | 1 |
| *Iberis amara* | 1 |
| *Lathyrus tuberosus* | 1 |
| *Lobelia erinus* | 1 |
| *Lotus corniculatus* | 1 |
| *Lysimachia punctata* | 1 |
| *Malus (apple)* | 1 |
| *Matricaria chamomilla* | 1 |
| *Melilotus officinalis* | 1 |
| *Mercurialis annua* | 1 |
| *Oenothera biennis* | 1 |
| *Oxalis corniculata* | 1 |
| *Papaver nudicaule* | 1 |
| *Phaseolus vulgaris* | 1 |
| *Pyracantha spp.* | 1 |
| *Rheum spp.* | 1 |
| *Ribes rubrum* | 1 |
| *Rosa multiflora* | 1 |
| *Rubus spp. (logan)* | 1 |
| *Rumex obtusifolius* | 1 |
| *Salvia officinalis* | 1 |
| *Symphoricarpos albus* | 1 |
| *Tanacetum parthenium* | 1 |
| *Valeriana officinalis* | 1 |
| *Valerianella locusta* | 1 |
| *Veronica serpyllifolia* | 1 |
| *Vicia cracca* | 1 |
| *Vinca major* | 1 |

**Table S8:** Linear mixed effect model output. Type II Analysis of Variance Table with Satterthwaite's method testing how urbanisation, area of cultivated flowers and supplement treatments (including wildflower additions, bee hotel additions and control sites), effects the abundance and species richness of bees (social and solitary), moths and hoverflies, and the seed set of tomatoes.

| Response | Terms | Coefficients | Estimate | Std. Error | df | t value | p value |
|---|---|---|---|---|---|---|---|
| Insect abundance | | | | | | | |
| | Treatment | | | | | | |
| | | Treatment – Control (intercept) | 2.774e+01 | 6.795e+00 | 3.877e+01 | 4.082 | 0.000216 |
| | | Treatment - Trap nest addition | -5.471e+00 | 4.605e+00 | 7.751e+01 | -1.188 | 0.238435 |
| | | Treatment - Flower addition | -5.099e+00 | 3.913e+00 | 7.610e+01 | -1.303 | 0.196478 |
| | Insect taxa | | | | | | |
| | | Moths (intercept) | 2.774e+01 | 6.795e+00 | 3.877e+01 | 4.082 | **0.000216** |
| | | Social | 8.505e+00 | 4.889e+00 | 7.395e+01 | 1.740 | 0.086070 |
| | | Solitary | -2.050e+01 | 5.071e+00 | 7.431e+01 | -4.043 | **0.000128** |
| | | Hoverfly | -1.633e+01 | 4.889e+00 | 7.395e+01 | -3.340 | **0.001314** |
| | Urbanisation | | 2.472e-06 | 3.548e-05 | 2.662e+01 | 0.070 | 0.944968 |
| Insect species richness | | | | | | | |
| | Treatment | | | | | | |
| | | Treatment – Control (intercept) | 1.111e+01 | 1.298e+00 | 4.123e+01 | 8.557 | **1.10e-10** |
| | | Treatment - Trap nest addition | -2.001e+00 | 8.539e-01 | 7.733e+01 | -2.343 | **0.0217** |
| | | Treatment - Flower addition | -4.170e-01 | 7.246e-01 | 7.575e+01 | -0.575 | 0.5667 |
| | Insect taxa | | | | | | |
| | | Moths (intercept) | 1.111e+01 | 1.298e+00 | 4.123e+01 | 8.557 | **1.10e-10** |
| | | Social | -7.169e+00 | 9.041e-01 | 7.402e+01 | -7.929 | **1.76e-11** |
| | | Solitary | -7.402e+00 | 9.381e-01 | 7.432e+01 | -7.891 | **2.03e-11** |
| | | Hoverfly | -5.794e+00 | 9.041e-01 | 7.402e+01 | -6.408 | **1.22e-08** |
| | Urbanisation | | -1.832e-06 | 6.835e-06 | 3.194e+01 | -0.268 | 0.7904 |
| Mean tomatoes seeds (log) | | | | | | | |
| | Treatment | | | | | | |
| | | Treatment – Control (intercept) | -0.47985 | 1.39486 | 19.91797 | -0.344 | 0.7344 |
| | | Treatment - Trap nest addition | 0.02276 | 0.09499 | 14.22874 | 0.240 | 0.8141 |
| | | Treatment - Flower addition | 0.23479 | 0.08723 | 13.57328 | 2.692 | **0.0179** |
| | Area of cultivated flowers | | 1.444e-03 | 4.482e-04 | 1.194e+02 | 3.222 | **0.00164** |

**Table S9:** Linear mixed effect model output. Type II Analysis of Variance Table with Satterthwaite's method testing the abundance of bees (social and solitary), moths and hoverflies, and if the mean abundance differs across habitat supplement treatments, and how urbanisation influences insect abundance.

| | Insect abundance | | |
|---|---|---|---|
| *Predictors* | *Estimates* | *CI* | *p* |
| (Intercept) | 6.01 | -110.97 – 122.99 | 0.919 |
| Insect taxa [Moth] | 20.49 | 8.71 – 32.27 | **0.001** |
| Insect taxa [Social bee] | 58.66 | 47.13 – 70.18 | **<0.001** |
| Insect taxa [Solitary bee] | 7.36 | -4.16 – 18.89 | 0.209 |
| Treatment [Flower addition] | 2.18 | -4.10 – 8.47 | 0.494 |
| Time [Timpoint 2] | 13.74 | 2.22 – 25.26 | **0.020** |
| Area of impervious surface [log] | -0.30 | -10.12 – 9.52 | 0.952 |
| social [Moth] × Time [timepoint 2] | -11.27 | -28.32 – 5.78 | 0.194 |
| Insect [Social bee] × Time [timepoint 2] | 7.51 | -8.69 – 23.72 | 0.362 |
| social [Solitary bee] × Time [timepoint 2] | -17.51 | -33.80 – -1.21 | **0.035** |
| **Random Effects** | | | |
| $\sigma^2$ | 399.88 | | |
| $\tau_{00}$ Block | 31.06 | | |
| ICC | 0.07 | | |
| N Block | 8 | | |
| Observations | 182 | | |
| Marginal $R^2$ / Conditional $R^2$ | 0.634 / 0.661 | | |

**Table S10:** Linear model output. Type II Analysis of Variance Table with Satterthwaite's method testing the species richness of bees (social and solitary), moths and hoverflies, and if the mean species richness differs across habitat supplement treatments, and how urbanisation influences insect species richness, and post hoc results of significant urbanisation x insect taxa interaction.

| | Insect species richness | | |
|---|---|---|---|
| *Predictors* | *Estimates* | *CI* | *p* |
| (Intercept) | 2.69 | -27.26 – 32.65 | 0.859 |
| Insect taxa [Moth] | 62.20 | 25.06 – 99.35 | **0.001** |
| Insect taxa [Social] | 11.82 | -24.90 – 48.54 | 0.526 |
| Insect taxa [Solitary] | -11.45 | -48.18 – 25.28 | 0.539 |
| Treatment [Flowers] | 0.09 | -0.82 – 0.99 | 0.853 |
| Time [Timpoint 2] | 4.42 | 2.78 – 6.06 | **<0.001** |
| Area of impervious surface [log] | -0.07 | -2.63 – 2.49 | 0.959 |
| Insect taxa [Moth] × Time [Timpoint 2] | -4.93 | -7.35 – -2.51 | **<0.001** |
| Insect taxa [Social] × Time [Timpoint 2] | -5.92 | -8.22 – -3.62 | **<0.001** |
| Insect taxa [Solitary] × Time [Timpoint 2] | -4.75 | -7.06 – -2.43 | **<0.001** |
| Insect taxa [Moth] × Area of impervious surface [log] | -4.44 | -7.55 – -1.32 | **0.006** |
| Insect taxa [Social] × Area of impervious surface [log] | -0.67 | -3.75 – 2.41 | 0.668 |
| Insect taxa [Solitary] × Area of impervious surface [log] | 1.10 | -1.98 – 4.18 | 0.482 |
| Observations | 182 | | |
| $R^2$ / $R^2$ adjusted | 0.606 / 0.559 | | |

Urbanisation

Area impervious surface * Insect taxa

| | | df | F ratio | p-value |
|---|---|---|---|---|
| *Post hoc:* | Hoverfly | 1 | 0.031 | 0.85 |
| | Moth | 1 | 11.71 | **0.0008** |
| | Social | 1 | 0.57 | 0.57 |
| | Solitary | 1 | 0.64 | 0.43 |

**Table S11:** Linear mixed effect model output. Type II Analysis of Variance Table with Satterthwaite's method testing the abundance of bees (social and solitary), moths and hoverflies, and if the mean abundance differs across habitat supplement treatments, and how the area of cultivated flowers influences insect abundance. Post hoc test reported when there was a significant cultivated flowers x insect taxa interaction.

| | Insect abundance | | |
|---|---|---|---|
| *Predictors* | *Estimates* | *CI* | *p* |
| (Intercept) | -2.91 | -54.37 – 48.56 | 0.911 |
| Insect taxa [Moth] | 30.05 | -44.05 – 104.15 | 0.424 |
| Insect taxa [Social] | -67.85 | -139.18 – 3.49 | 0.062 |
| Insect taxa [Solitary] | 5.51 | -65.85 – 76.86 | 0.879 |
| Treatment [Flowers] | 0.59 | -5.47 – 6.65 | 0.848 |
| Time [Timpoint 2] | 14.11 | 2.87 – 25.35 | **0.014** |
| Area Cultivated flowers [log] | 0.92 | -7.14 – 8.98 | 0.822 |
| Insect taxa [Moth] × Time [Timpoint 2] | -11.50 | -27.97 – 4.98 | 0.170 |
| Insect taxa [Social] × Time[Timpoint 2] | 7.19 | -8.61 – 23.00 | 0.370 |
| Insect taxa [Solitary] × Time [Timpoint 2] | -17.36 | -33.26 – -1.47 | **0.032** |
| Insect taxa [Moth] × Area Cultivated flowers [log] | -1.49 | -13.09 – 10.11 | 0.800 |
| Insect taxa [Social] × Area Cultivated flowers [log] | 20.11 | 8.97 – 31.25 | **<0.001** |
| social [Solitary] × Area Cultivated flowers [log] | 0.24 | -10.90 – 11.38 | 0.966 |
| **Random Effects** | | | |
| $\sigma^2$ | 363.86 | | |
| $\tau_{00\ Block}$ | 12.60 | | |
| ICC | 0.03 | | |
| $N_{Block}$ | 8 | | |
| Observations | 176 | | |
| Marginal $R^2$ / Conditional $R^2$ | 0.685 / 0.696 | | |

Area cultivated flowers

Area cultivated flowers * Insect taxa

| | | df | F-ratio | p-value |
|---|---|---|---|---|
| *Post hoc:* | Hoverfly | 1 | 0.05 | 0.824 |
| | Moth | 1 | 0.07 | 0.90 |
| | Social | 1 | 26.17 | **<0.001** |
| | Solitary | 1 | 0.079 | 0.78 |

**Table S12:** Linear model output. Type II Analysis of Variance Table with Satterthwaite's method testing the species richness of bees (social and solitary), moths and hoverflies, and if the mean species richness differs across habitat supplement treatments, and how the area of cultivated flowers influences insect species richness.

| | Insect species richness | | |
|---|---|---|---|
| *Predictors* | *Estimates* | *CI* | *p* |
| (Intercept) | 1.84 | -3.13 – 6.81 | 0.465 |
| Insect taxa [Moth] | 9.41 | 7.63 – 11.18 | **<0.001** |
| Insect taxa [Social] | 3.76 | 2.00 – 5.52 | **<0.001** |
| Insect taxa [Solitary] | 1.59 | -0.17 – 3.35 | 0.076 |
| Treatment [Flowers] | 0.00 | -0.98 – 0.98 | 0.998 |
| Time [Timepoint 2] | 4.50 | 2.74 – 6.26 | **<0.001** |
| Area of Culitvaed Flowers [log] | 0.09 | -0.66 – 0.84 | 0.808 |
| Insect taxa [Moth] × Time [Timepoint 2] | -5.08 | -7.65 – -2.51 | **<0.001** |
| Insect taxa [Social] × Time[Timepoint 2] | -5.89 | -8.36 – -3.42 | **<0.001** |
| Insect taxa [Solitary] × Time[Timepoint 2] | -4.76 | -7.25 – -2.28 | **<0.001** |
| Observations | 176 | | |
| $R^2$ / $R^2$ adjusted | 0.567 / 0.523 | | |

**Table S13:** Linear model output. Type II Analysis of Variance Table with Satterthwaite's method testing the estimate Shannon diversity- derived from the abundance-based rarefaction- of bees (social and solitary), moths and hoverflies, and if the mean estimated Shannon diversity differs across habitat supplement treatments, and how the area of cultivated flowers influences insect Shannon diversity.

| | **Estimated Shannon Diversity** | | |
|---|---|---|---|
| *Predictors* | *Estimates* | *CI* | *p* |
| (Intercept) | 5.62 | 1.31 – 9.93 | **0.011** |
| Time [Timepoint 2] | 2.95 | 0.76 – 5.13 | **0.009** |
| Treatment [Flowers] | -0.07 | -0.84 – 0.69 | 0.850 |
| Area of Cultivated Flowers [log] | -0.39 | -0.98 – 0.20 | 0.193 |
| Insect taxa [Moth] | 9.38 | 6.58 – 12.19 | **<0.001** |
| Insect taxa [Social] | 1.07 | -0.75 – 2.90 | 0.247 |
| Insect taxa [Solitary] | 0.03 | -2.07 – 2.12 | 0.980 |
| Time [Timpoint 2] × Insect taxa [Moth] | -8.12 | -11.87 – -4.36 | **<0.001** |
| Time [Timepoint 2] × Insect taxa [Social] | -4.54 | -6.90 – -2.19 | **<0.001** |
| Time [Timepoint 2] × Insect taxa [Solitary] | -3.02 | -5.80 – -0.24 | **0.033** |
| Observations | 158 | | |
| $R^2$ / $R^2$ adjusted | 0.442 / 0.379 | | |

**Table S14**: Linear model output. Type II Analysis of Variance Table with Satterthwaite's method testing the estimate Shannon diversity- derived from the abundance-based rarefaction- of bees (social and solitary), moths and hoverflies, and if the mean estimated Shannon diversity differs across habitat supplement treatments, and how the area of impervious surfaces effect insect Shannon diversity. Post hoc test reported when there was a significant urbanisation x insect taxa interaction.

| | **Estimated Shannon Diversity** | | | |
|---|---|---|---|---|
| *Predictors* | *Estimates* | *CI* | *p* | |
| (Intercept) | 9.79 | -23.70 – 43.29 | 0.564 | |
| Time [Timepoint 2] | 2.93 | 0.86 – 5.01 | **0.006** | |
| Treatment [Flowers] | -0.01 | -0.74 – 0.72 | 0.984 | |
| Insect taxa [Moth] | 48.19 | -7.38 – 103.75 | 0.089 | |
| Insect taxa [Social] | 0.94 | -33.63 – 35.51 | 0.957 | |
| Insect taxa [Solitary] | -36.16 | -78.32 – 5.99 | 0.092 | |
| Time [Timpoint 2] × Insect taxa [Moth] | -7.98 | -11.59 – -4.37 | **<0.001** | |
| Time [Timepoint 2] × Insect taxa [Social] | -4.63 | -6.86 – -2.39 | **<0.001** | |
| Time [Timepoint 2] × Insect taxa [Solitary] | -1.92 | -4.72 – 0.88 | 0.177 | |
| Area of impervious surfaces [log] | -0.57 | -3.38 – 2.25 | 0.692 | |
| Area of impervious surfaces [log] × Insect taxa [Moth] | -3.25 | -7.89 – 1.40 | 0.169 | |
| MM 250m [log] × Insect taxa [Social] | 0.02 | -2.87 – 2.90 | 0.991 | |
| Area of impervious surfaces [log] × Insect taxa [Solitary] | 3.03 | -0.49 – 6.54 | 0.091 | |
| Observations | 158 | | | 164 |
| $R^2$ / $R^2$ adjusted | 0.442 / 0.379 | | | 0.472 / 0.403 |

Area of impervious surface

| | Area cultivated flowers * Insect taxa | df | F-ratio | p-value |
|---|---|---|---|---|
| *Post hoc:* | Hoverfly | 1 | 0.16 | 0.70 |
| | Moth | 1 | 3.59 | 0.60 |
| | Social | 1 | 0.49 | 0.48 |
| | Solitary | 1 | 3.7 | **0.056** |

**Table S15:** Linear mixed effect model output. Type II Analysis of Variance Table with Satterthwaite's method testing if the mean tomato seeds differ across habitat supplement treatments, and how the area of urbanisation influences number of tomato seeds.

| Response | Terms | Coefficients | Estimate | Std. Error | df | t value | p value |
|---|---|---|---|---|---|---|---|
| log(Tomato seeds) | | | | | | | |
| | Treatment | | | | | | |
| | | Treatment – Control (intercept) | -0.53055 | 1.34449 | 20.80029 | -0.395 | 0.69715 |
| | | Treatment - Flower addition | 0.22532 | 0.07397 | 14.85742 | 3.046 | **0.00824** |
| | Urbanisation | | | | | | |
| | | | 0.33885 | 0.11278 | 20.77962 | 3.005 | **0.00680** |

**Table S16:** Linear mixed effect model output. Type II Analysis of Variance Table with Satterthwaite's method testing if the mean tomato seeds differ across habitat supplement treatments, and how the area of cultivated flowers influences number of tomato seeds.

| Response | Terms | Coefficients | Estimate | Std. Error | df | t value | p value |
|---|---|---|---|---|---|---|---|
| log(Tomato seeds) | | | | | | | |
| | Treatment | | | | | | |
| | | Treatment – Control (intercept) | 2.76790 | 0.32929 | 18.08671 | 8.406 | 1.16e-07 |
| | | Treatment - Flower addition | 0.20427 | 0.07852 | 14.40859 | 2.601 | **0.0205** |
| | Area of cultivated flowers | | | | | | |
| | | | 0.11740 | 0.05123 | 17.19082 | 2.291 | **0.0348** |

**Table S17:** Linear mixed effect model output. Type II Analysis of Variance Table with Satterthwaite's method testing how species richness of bees influences number of tomato seeds.

| Response | Terms | Coefficients | Estimate | Std. Error | df | t value | p value |
|---|---|---|---|---|---|---|---|
| log(Tomato seeds) | | | | | | | |
| | Bee species richness (intercept) | | 29.4723 | 4.1676 | 32.1916 | 7.072 | **4.92e-08** |
| | | | 1.3441 | 0.6102 | 43.6256 | 2.203 | **0.0329** |

**Table S18**: Linear mixed effect model outputs. Type II Analysis of Variance Table with Satterthwaite's method testing if network structure of bees (social, solitary) and hoverflies differ across habitat supplement treatments, and how the area of cultivated flowers influences these network metrics (total number of plant species, linkage density, generality of insects).

| Response | Terms | Coefficients | Estimate | Std. Error | df | t value | p value |
|---|---|---|---|---|---|---|---|
| Total number of plant species | | | | | | | |
| | Treatment | | | | | | |
| | Treatment – Control (intercept) | | 3.6503 | 2.0231 | 112.6090 | 1.804 | 0.07385 |
| | Treatment - Flower addition | | 0.3524 | 0.4868 | 134.4616 | 0.724 | 0.47029 |
| | Insect taxa | | | | | | |
| | | Social (Intercept) | 3.6503 | 2.0231 | 112.6090 | 1.804 | **0.07385** |
| | | Solitary | -6.9583 | 0.5730 | 132.3357 | -12.143 | **< 2e-16** |
| | | Hoverfly | -5.5625 | 0.5730 | 132.3357 | -9.707 | **< 2e-16** |
| | Area of cultivated flowers | | 1.444e-03 | 4.482e-04 | 1.194e+02 | 3.222 | **0.00164** |
| Linkage density | | | | | | | |
| | Treatment | | | | | | |
| | Treatment – Control (intercept) | | 2.556e+00 | 1.279e-01 | 1.390e+02 | 19.989 | **< 2e-16** |
| | Treatment - Flower addition | | 6.868e-02 | 1.148e-01 | 1.390e+02 | 0.598 | 0.55056 |
| | Insect taxa | | | | | | |
| | | Social (Intercept) | 2.556e+00 | 1.279e-01 | 1.390e+02 | 19.989 | **< 2e-16** |
| | | Solitary | -1.350e+00 | 1.352e-01 | 1.390e+02 | -9.988 | **< 2e-16** |
| | | Hoverfly | -1.078e+00 | 1.352e-01 | 1.390e+02 | -7.975 | **5.05e-13** |
| | Area of cultivated flowers | | 2.780e-04 | 9.752e-05 | 1.390e+02 | 2.851 | **0.00503** |
| Generality of insect (number of plants per insect species) | | | | | | | |
| | Treatment | | | | | | |
| | Treatment – Control (intercept) | | 1.9331 | 0.4859 | 139.0000 | 3.979 | **0.000111** |
| | Treatment - Flower addition | | 0.0161 | 0.1259 | 139.0000 | 0.128 | 0.898462 |
| | Insect taxa | | | | | | |
| | | Social (Intercept) | 1.9331 | 0.4859 | 139.0000 | 3.979 | **0.000111** |
| | | Solitary | -1.5660 | 0.1491 | 139.0000 | -10.506 | **< 2e-16** |
| | | Hoverfly | -1.3771 | 0.1491 | 139.0000 | -9.238 | **3.85e-16** |
| | Area of cultivated flowers | | 2.064e-04 | 1.076e-04 | 1.390e+02 | 1.919 | **0.057** |

### Supplementary Text:

**Text S1:** Flower patch addition methodology:

Site preparation (March):

- Cut vegetation down to 5-10cm and turn the soil over in an area of 100$m^2$
- Remove persistent weeds

Seed bed preparation and sowing (April-May):

- Rake over soil and remove debris and stones.
- Sow 3g/$m^2$ of seeds evening throughout the patch (on a calm, dry day).
- Roll over the seeds lightly to maximise germination.
